# The effects of group size and assortment on the evolution of division of labor

**DOI:** 10.64898/2026.05.22.724929

**Authors:** Aidan Fielding, Erol Akçay, Joshua Plotkin

## Abstract

Stable variation in public-good production can generate biological division of labor. Two key drivers are the size of interacting groups and the degree of assortment (relatedness) among individuals with similar investment levels. Here we extend adaptive-dynamic models of continuous public-goods investment by allowing assortment in group formation. We show that increasing group size typically enlarges the range of benefit and cost curvatures that permit evolutionary branching, whereas assortment tends to shrink this range and prevents branching under complete relatedness. For a broad class of models, branching requires diminishing marginal public benefits and diminishing marginal private costs, with costs decreasing faster than benefits. Finally, we analyze post-branching dynamics for one and two public goods, and find that assortment can stabilize the resulting two-type division of labor. Together, these results show how group size, assortment, and payoff curvature jointly determine when heterogeneity in public-goods production can evolve.

## 1 Background

Variation in the production of public goods occurs in a wide range of taxa, from microbes to humans. In bacterial populations, producers and non-producers of iron-scavenging siderophores coexist across diverse environments [Cordero et al., 2012; Kramer et al., 2019; Ross-Gillespie et al., 2014; Schiessl et al., 2019], and similar within-colony variation occurs for other public goods such as proteases and biofilm substrates [Drescher et al., 2014; Smith and Schuster, 2019; van Gestel et al., 2015]. In many systems, individuals invest in more than one public good, and variation in these investments can lead to task specialization and division of labor. For example, mound-building mice differ in whether they transport vegetation or roof-building materials during nest construction [Hurtado et al., 2013; Smith and Riehl, 2022]. Human hunter-gatherers vary in their rates of meat versus plant foraging [Anderson et al., 2023; Apicella and Silk, 2019; Stibbard-Hawkes et al., 2022]. Even among cancer cells, individuals differ in acid production and vascularization, two traits that jointly facilitate metastasis [Kaznatcheev et al., 2017]. Across these systems, individuals allocate effort differently across public goods, resulting in functional specialization within a shared ecological context.

Two key ecological parameters shape the production of public goods: group size and relatedness among interacting individuals [Archetti and Scheuring, 2012; Frank, 2010; Hamilton, 1964; Motro, 1991; Netz et al., 2025; Rankin et al., 2007]. The example of iron-scavenging bacteria illustrates both. Individual cells secrete siderophores that diffuse into the environment, bind to iron, and facilitate uptake by nearby cells [Bergeron et al., 1985; Bergeron and Weimar, 1990]. When diffusion is localized, only a few neighboring cells benefit from a focal cell’s secretion, so the per-capita benefit remains high. When diffusion is extensive, many cells benefit, and the per-capita return to the producer declines. Larger effective group size therefore tends to reduce the incentive to produce the public good. At the same time, larger groups may permit greater variation: a single free-rider has a smaller effect on the total public benefit. Relatedness adds a second layer. A focal cell typically shares lineage with some fraction of its neighbors [Cordero et al., 2012]. Higher relatedness increases the inclusive fitness return to producing siderophores, because benefits accrue to genetic relatives. But greater relatedness can also reduce phenotypic variation, since relatives are more similar. Taken together, group size and relatedness exert opposing and interacting effects on both the level of public-good production and the emergence of division of labor. Their joint influence is therefore not obvious a priori. A theoretical framework that disentangles the effects of group size and relatedness is therefore needed. Such a framework should identify ecologically meaningful conditions under which multiple public-good production levels can emerge and coexist, leading to division of labor. It should also clarify how changes in group size or relatedness shift those conditions.

The conditions for division of labor depend not only on group size and relatedness [Wakano and Lehmann, 2014], but also on how the benefits and costs to individuals change as a function of investments. The public benefit depends jointly on all group members’ investments, whereas the private cost depends only on the focal individual’s investment. Like group size and relatedness, these functions have direct ecological meaning. In iron-scavenging bacteria, for example, the rate of iron uptake for a focal cell is an increasing function of its own secretion of siderophores and that of its neighbors, as well as the concentration of iron in the extracellular environment [Bergeron and Weimar, 1990; Kramer et al., 2019]. At the same time, producing siderophores imposes a private cost on the focal cell due to both the deterioration of cellular machinery and the opportunity cost of devoting resources to iron-scavenging instead of other essential metabolic processes. Crucially, evolutionary outcomes depend not just on the levels of benefit and cost, but on how they change with investment—that is, on the curvatures of *B* and *C*. These curvatures represent ecological constraints that are, in principle, measurable properties of real systems.

To disentangle these effects, we develop a minimal theoretical framework. We assume that the population is initially monomorphic, so any polymorphism must arise through evolutionary diversification, and that there is no class structure or asymmetry that predisposes individuals to particular phenotypes. These assumptions isolate the role of ecological parameters—group size and relatedness—in generating division of labor. We model an individual’s level of public-good production as a continuously varying quantitative trait, which we call an “investment.” Continuous variation allows evolution through small mutational steps, without requiring large phenotypic jumps. In this setting, polymorphism can arise via evolutionary branching: a monomorphic population evolves to a critical trait value under directional selection, and disruptive selection near that point causes the population to split into two types that subsequently diverge [Dieckmann and Law, 1996; Doebeli and Dieckmann, 2000; Geritz et al., 1998; Leimar, 2009; Metz et al., 1996]. Evolutionary branching thus provides a general mechanism by which division of labor on public goods can emerge from an initially homogeneous population [Doebeli et al., 2004; Fielding et al., 2025; Henriques et al., 2021; Lehmann and Mullon, 2025].

Evolutionary branching requires that the curvatures of the public benefit and private cost functions lie within a specific range, and this range depends on group size and relatedness. Previous studies have examined branching of investments in public goods without relatedness [Doebeli et al., 2004; Henriques et al., 2021]; others studies addressed branching of arbitrary traits with relatedness, but did not derive the general constraints on the benefit and cost functions that are required for branching [Lehmann and Mullon, 2025; Wakano and Lehmann, 2014]. A unified analysis that connects relatedness directly to the ecological structure of public-good benefits and costs has been lacking.

Here we incorporate relatedness into the analysis of evolutionary branching of investments in public goods. We show that the range of curvatures of the benefit and cost functions that leads to branching tends to grow with group size and shrink with increasing relatedness. In tandem, we show that branching requires the benefit and cost functions to diminish with respect to investments under much more general assumptions than those discussed in previous literature. Finally, we introduce relatedness into the analysis of polymorphic investments after branching, with both one and two public goods, and we find one intriguing difference from prior results: with positive relatedness, it is possible for the two-type division of labor after branching to be evolutionarily stable, meaning it has no potential for further diversification.

## 2 Methods

We consider a large population of individuals playing a continuous public goods game in groups of size *n* ≥ 2.

Focal individual *i* ∈ (1, …, *n*) invests amount *z*_i_ into the public good. The payoff that *i* receives from the game is

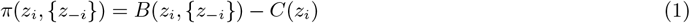

where {*z*_−i_}is the set of investments of the *n* − 1 non-focal players. The public benefit *B*(·) is a function of all *n* investments, and the private cost *C*(·) to *i* is a function of *i*’s investment only. Both *B*(·) and *C*(·) are real-valued, non-negative functions. A natural assumption is that investments increase both public benefits and private costs; that is, 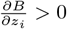and 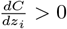for any *i* ∈ (1, …, *n*). We also assume that investments in the public good are interchangeable, so that if the investment levels of any two individuals are switched, their private costs switch, while the public benefit stays the same. This will be true when when there are no inherent differences between individuals other than (possibly) their investment levels.

A relevant alternative form of public goods payoff is the multiplicative form *π*(*z*_i_, {*z*_−i_*}*) = *B*(*z*_i_, {*z*_−i_*}*) · *C*(*z*_i_), where 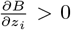and 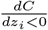for all *i*. This form is appropriate when benefit, cost and resulting payoff are defined as rates. It is used in standard analyses of reproductive division of labor, in which viability corresponds to public benefit and fecundity to private cost [Michod, 2006, 2007; Yanni et al., 2020]. As a robustness check, we repeated the analysis below with the multiplicative form of payoff and compared general results.

Public-good groups of size *n* are formed by a mixture of assortment between types and random association. We model assortment with own type by introducing a parameter *r* ∈ [0, 1], which gives the probability that each non-focal group mate *j* ≠ *i* will be of the same type as the focal player and therefore have the same trait value (i.e. *z*_*j*_ = *z*_i_); with probability 1 − *r*, the non-focal individual is drawn randomly from the population (this can yield the same type as the focal player but also other types).

The fitness of a focal individual *i* is equal to the expected payoff 𝔼 [*π*(*z*_i_, {*z*_−i_*}*)] that *i* receives from the game, given the group-formation mechanism described above. Thus, the expectation is taken over all possible group compositions within which *i* may play the public goods game. The probability of each composition depends on *i* due to assortment.

To derive the adaptive dynamics of investments, we derived the invasion fitness *w*_inv_ of a rare mutant type with investment *z*_mut_ in a monomorphic population with resident type *z*_res_ [Dieckmann and Law, 1996; Metz et al., 1996]. We denote by {*z}*_*k*+1_ a set of *n* investments of which *k* + 1 elements are equal to *z*_mut_ and *n* − 1 − *k* are equal to *z*_res_ (note that our interchangeability assumption means the ordering of the investments does not matter.) Because the mutant is initially rare, a focal mutant’s groupmates will have trait *z*_mut_ only with the assortment probability *r*. Thus, the probability distribution of the investment set {*z}*_*k*+1_ in a rare mutant scenario is going to be binomial with parameters *n* − 1 and *r*. Consequently, we can write the fitness *w*_inv_ of a mutant type in a monomorphic resident population:

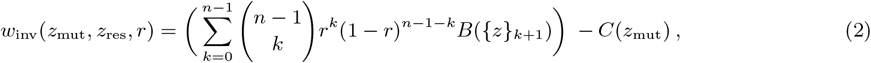

where the first term is the expected public benefit received by a mutant, and the second term is the private cost paid by a mutant. Under adaptive dynamics, the trait value will evolve in the direction of increasing invasion fitness *w*_inv_ [Dieckmann and Law, 1996]. i.e.,

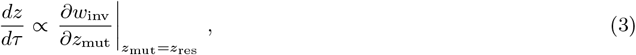

where *τ* denotes time at the evolutionary scale. When the partial derivative on the right hand side vanishes, evolutionary dynamics are at a critical point.

A critical trait value *z*_*_ is convergently stable if a population in which all residents invest *z* ≠ *z*_*_ (in an appropriate neighborhood of *z*_*_) will evolve to a state in which all residents invest *z*_*_. The critical value is evolutionarily unstable if a population in which all residents invest *z*_*_ can be invaded by mutants who invest slightly above or below *z*_*_. If *z*_*_ has both of these properties, evolutionary branching occurs: investments in a monomorphic population evolve to *z*_*_; the monomorphic residents are invaded by mutants with nearby trait values; the resident and invader types coexist at stable frequencies *p* ∈ (0, 1) and (1 − *p*), forming a dimorphic resident population; and this process iterates, causing the dimorphic types to evolve away from each other in trait space [Geritz et al., 1998].

More explicitly, these two properties can be expressed in terms of derivatives of mutant invasion fitness *w*_inv_. A critical trait value *z*_*_ is convergently stable and evolutionarily unstable if *w*_inv_ satisfies the following second-order conditions [Geritz et al., 1998; Leimar, 2009]:

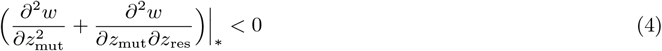

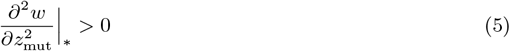

where “*” indicates that the derivatives are evaluated such that *z*_mut_ = *z*_res_ = *z*_*_. Note that for (5) and (4) to both be true, it is necessary that 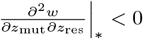. This is a general requirement for evolutionary branching [Lehmann and Mullon, 2025].

We expand the conditions for evolutionary branching in terms of the first and second derivatives of *B* and *C*, evaluated at a critical point where all investments are equal to *z*\*. These quantities are equal when taken with respect to any of the *n* investments (or any pair of investments, in the case of the cross derivative of *B*). Therefore the analysis simplifies to five quantities. For any two players *i* and *j* from 1 to *n*, where *i* ≠ *j*, the five quantities are defined in Table 1. The conditions for branching are expressed in terms of *n, r, B*_•,•_′, *B*_•,•_, and *C*_•,•_. Note that the latter three terms are functions of *n* and *r*, because they are evaluated at critical investment *z*_*_.

**Table 1:** First- and second-order effects of investments on public benefit and private cost.

|  |  |
| --- | --- |
| $B_{\bullet} = \frac{\partial B}{\partial z_i} \Big\vert _*$ | marginal public benefit of focal individual’s investment |
| $C_{\bullet} = \frac{dC}{dz_i} \Big\vert _*$ | marginal private cost of focal individual’s investment |
| $B_{\bullet, \bullet} = \frac{\partial^2 B}{\partial z_i^2} \Big\vert _*$ | rate of change of the marginal public benefit of focal individual’s investment |
| $C_{\bullet, \bullet} = \frac{d^2 C}{dz_i^2} \Big\vert _*$ | rate of change of the marginal private cost of a focal individual’s investment |
| $B_{\bullet, \bullet'} = \frac{\partial^2 B}{\partial z_i \partial z_j} \Big\vert _*$ | rate of change of the marginal public benefit of a coplayer’s investment |

In the results section below, we analyze what characteristics of *B*_•,•_′, *B*_•,•_, and *C*_•,•_ are required for branching to occur. Then, we show how changes in group size *n* and relatedness *r* affect branching conditions. To do this, we assume specific functional forms for public benefit *B* and private cost *C*. Results below focus on the setting in which public benefit is a function of the sum of investments, and benefit and cost are both power functions, that is

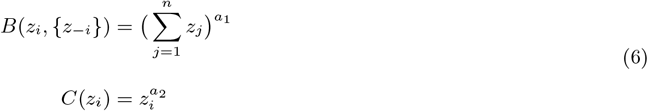

where *a*_1_ > 0 and *a*_2_ > 0, under the assumption that benefit and cost both increase with investments. This form of payoff using power functions has minimal terms and parameters, allowing an intuitive explanation of how each individual’s investment is functionally converted into public benefit *B* and private cost *C*: an investment into *B* is added to a sum whose total benefit either decelerates or accelerates with size; the cost *C* of that investment also either decelerates or accelerates with its size. The concavity (or convexity) of public benefit and private cost are governed by the parameters *a*_1_ and *a*_2_.

As a robustness check, we carried out the analysis assuming that public benefit and private cost are quadratic functions, that is

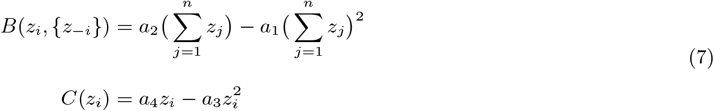

where all *a*s are positive. These functions are both increasing in the focal investment, *z*_i_ for some range of positive *z*_i_, but will start decreasing for high enough focal investments. To avoid benefits or costs decreasing with investment, an upper limit is imposed on the investments at the maximum of *B* or *C* (whichever is lower; see S.I. B.8.2 for more details). This quadratic payoff function and constraints were used in previous studies of the adaptive dynamics of investments in a public good [Doebeli et al., 2004; Henriques et al., 2021].

### 2.1 Adaptive dynamics after branching

We extend our analysis to the adaptive dynamics of dimorphic investments after evolutionary branching. A dimorphic population has two types, *u* and *v*, with investments *z*_*u*_ and *z*_*v*_, and frequencies *p* ∈ [0, 1] and 1 − *p*. With two resident types, the group composition of a rare mutant has a multinomial distribution: with probability *r*, a given non-focal player will share the mutant’s trait; with probability (1 − *r*)*p*, then non-focal player will be type *u*; and with probability (1 − *r*)(1 − *p*), the non-focal player will be type *v*. For payoff models 1 and 2, we check for the existence of critical points (*z*_*u*_, *z*_*v*_)* and determine the convergence and evolutionary (in)stability of any such points. These properties are computed analogously to the monomorphic case (S.I. B.9). We are particularly interested in whether there are any qualitative differences between dynamics with no assortment (*r* = 0) and those with partial assortment (0 < *r* < 1).

### 2.2 Two public goods

Finally, we extend our analysis to the case with two public goods. The simplest way to define fitness in this setting is to add the payoffs of the two public goods together. This leads to a payoff function of the form *π* = *B*_1_ −*C*_1_ +*B*_2_ −*C*_2_, where *B*_1_ (*B*_2_) and *C*_1_ (*C*_2_) are the public benefit and private cost functions for good 1 (good 2), respectively. A further simplifying assumption is that the public benefit and private cost of good *k* are functions only of investments in good *k*. Thus, there are no synergistic or antagonistic interactions between the two goods. The adaptive dynamics of investments in this setting with no assortment (*r* = 0) have been studied [Fielding et al., 2025; Henriques et al., 2021]. We want to know if there are any qualitative differences between dynamics with no assortment and those with partial assortment.

## 3 Results

### 3.1 Selection gradient

The selection gradient with assortment *r* is

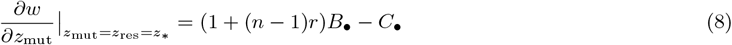

This gives the direction of selection on investments, which for a one-dimensional trait reduces to up- or downward selection. At a critical investment level *z*_*_, the selection gradient vanishes, i.e., (1 + (*n* − 1)*r*)*B*_•_ − *C*_•_ = 0. This is a version of Hamilton’s rule in marginal form, where direct fitness effects are given by *B*_•_ − *C*_•_ and indirect fitness effects via assortment are given by *r*(*n* − 1)*B*_•_ [Lehmann and Mullon, 2025].

The critical investment *z*_*_ increases with *r*, that is 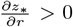, if (1 + (*n* − 1)*r*)(*B*_•,•_ + (*n* − 1)*B*_•,•_′) < *C*_•,•_ (S.I. B.6). The latter inequality also implies that *z*_*_ is convergence stable, as explained in the next section.

### 3.2 Evolutionary branching conditions with assortment

For investments to undergo evolutionary branching, the critical investment *z*_*_ has to be convergence stable but evolutionary unstable. These two conditions are met if and only if the following inequalities are satisfied:

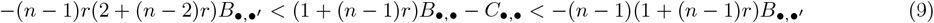

The right inequality in (9) is required for *z*_*_ to be convergent stable, while the left inequality is required for *z*_*_ to be evolutionarily unstable (S.I. B.1 and B.2). The middle term (1 + (*n* − 1)*r*)*B*_•,•_ − *C*_•,•_ is the second partial derivative of fitness with respect to a focal investment, which indicates how a small change in a focal investment will change the marginal inclusive fitness due to additional focal investment. The right-hand term is the negative of the sum across all nonfocal players of the cross derivatives of the inclusive fitness benefit with respect to a focal and a nonfocal investments. This indicates how a small change in a focal investment will change the marginal inclusive fitness benefit due to a small change in a related nonfocal player’s investment, for all coplayers. The convergence stability condition thus requires that the second-order fitness effect of a focal investment on itself is less than the negative sum of the second-order fitness effects of the focal investment on all coplayers, scaled by the probability *r* with which coplayers share the focal’s type. The ecological intuition for this is that individuals are weighing the second-order fitness effect of their investment on themselves against the summed second-order effects on related coplayers. The second-order effect with respect to one coplayer is *B*_•,•_′. Consider the case where *B*_•,•_′ is negative. Then the right-hand term in (9) will be positive, and the critical point can retain convergence stability even if focal fitness accelerates (i.e. the middle term is positive), because the negative second order effect of focal investment on related coplayers provides a disincentive for the focal to invest that eventually balances the individual incentive due to acceleration. Conversely, if *B*_•,•_′ is positive, the second order effect of focal investment on related coplayers is an incentive to invest more. In that case, the right-hand term in (9) is negative, and convergence stability is only possible if focal fitness decelerates (i.e. the middle term is negative).

The left-hand term in (9) is the negative of the sum across all nonfocal players of the cross derivatives of inclusive fitness benefit with respect to a nonfocal investment and the *n* − 2 other nonfocal investments. It denotes how a small change in a related coplayer’s investment will change the marginal inclusive fitness benefit due to a small change in another related coplayer’s investment, for all coplayers. Consider again the case where *B*_•,•_′ < 0. If a focal individual increases (decreases) their investment, the marginal inclusive public benefit decreases (increases) for all coplayers’ investments, reducing coplayers’ incentive to invest more (less). When one individual changes their investment (either up or down), the coplayers are further incentivized to not change their investment. If in addition, focal fitness is accelerating, then individuals have an incentive to change their investments away from those of their coplayers. The combination of these two incentives causes disruptive selection. As we show below, they are both required for evolutionary branching to occur.

#### 3.2.1 Public benefit and private cost functions that satisfy branching conditions

If there is zero or partial relatedness in the population (0 ≤ *r* < 1), the outer inequality in (9) requires that *B*_•,•_′ < 0 (S.I. B.3). This means a marginal increase in a focal player’s investment will decrease the marginal public benefit of a coplayer’s investment (or vice versa). The left inequality in (9) then implies that the middle term is positive, that is 0 < (1 + (*n* − 1)*r*)*B*_•,•_ − *C*_•,•_, meaning the inclusive fitness of a focal individual is accelerating with respect to their own investment in the public good. It follows that *C*_•,•_ < (1 + (*n* − 1)*r*)*B*_•,•_, so the change in the marginal private cost due to an increase in the focal individual’s investment is less than the change in the marginal inclusive public benefit. All three of these constraints on the benefit and cost functions are easy to see in the case of no relatedness (*r* = 0), in which inequalities (9) simplify to 0 < *B*_•,•_ − *C*_•,•_ < −(*n* − 1)*B*_•,•_′. The no-relatedness case is a generalization of a result from Doebeli et al. [2004], who assumed that public benefit is specifically a function of the sum of all group members’ investments.

If the public benefit is a function of the sum or product of investments, then its second partial derivative with respect to a focal player’s investment is less than or equal to the cross derivative with respect to the focal and a coplayer’s investment (i.e. *B*_•,•_ ≤ *B*_•,•_′). In fact, this inequality is true for a much broader family of public benefit functions, which we define in S.I. B.4. Branching conditions (9) in this setting therefore require that *B*_•,•_ < 0. In other words, the public benefit received by a focal individual is decelerating with respect to their own investment. It follows that *C*_•,•_ < (1 + (*n* − 1)*r*)*B*_•,•_ < 0, so the private cost paid by a focal individual is also decelerating with respect to their own investment. Thus, for a general class of plausible public benefit functions, evolutionary branching requires that the marginal private cost decreases faster than the marginal inclusive public benefit does with respect to a focal individual’s investment.

#### 3.2.2 Analysis with multiplicative payoff form

We derived evolutionary branching conditions similar to (9) for payoff functions of the form *π*(*z*_i_, {*z*_−i_*}*) = *B*(*z*_i_, {*z*_−i_*}*) · *C*(*z*_i_) for all *i* (S.I. D). As with the additive form, the branching conditions for this alternative form require that *B*_•,•_′ < 0, so a marginal increase in a coplayer’s investment must decrease the marginal public benefit of a focal player’s investment (or vice versa). Again, if public benefit is a function of the sum or product of investments, then branching requires that the benefit received by a focal individual is decelerating with respect to their own investment (i.e. *B*_•,•_ ≤ *B*_•,•_′ < 0). However, in this setting, there is no restriction on the concavity of the private cost function.

### 3.3 Effect of group size (*n*) and relatedness (*r*) on branching conditions

If there is complete relatedness in the population (*r* = 1), then the left and right terms in inequalities (9) are equal, so the branching conditions cannot be satisfied. This means that branching cannot occur for fully assorted groups. It is notable that *r* = 1 is the standard assumption in the theory of reproductive division of labor. In that setting, the existence of two types with distinct traits must be imposed exogenously [Cooper et al., 2022; Cooper and West, 2018; Rueffler et al., 2012; West and Cooper, 2016]. Our results here suggest that two types could not evolve incrementally from one ancestral type with full assortment.

With partial relatedness (0 < *r* < 1), if the monomorphic critical point *z*_*_ is convergence stable, then *z*_*_ increases with *r* (see section 3.1 and S.I. B.6). This is natural: as focal individuals interact more frequently with related coplayers, their inclusive fitness increases more, for a given investment in the public good. Because *B*_•,•_′, *B*_•,•_, and *C*_•,•_ are functions of *n* and *r*, analyzing how *n* and *r* change evolutionary branching conditions (9) requires additional assumptions about the functional forms of public benefit *B* and private cost *C*.

#### 3.3.1 Power-function payoff model

We applied the analysis for the example case of public benefit and private cost functions specified by power functions (equation (6); S.I. B.8.1). The effects of *n* and *r* can readily be interpreted in terms of parameters *a*_1_ and *a*_2_, which govern the concavity of benefit and cost, respectively.

The evolutionary branching conditions with power function payoffs simplify to

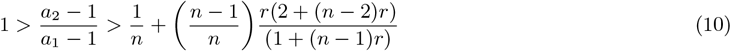

This requires that 0 < *a*_1_ < *a*_2_ < 1, so both benefits and costs must decelerate, as shown in section 3.2.1. The further requirement that marginal private benefit decrease faster than marginal inclusive public benefit (left inequality in (9)) is satisfied if *n*(*a*_2_ − 1) < *a*_1_ − 1. As *n* increases, the rightmost term in (10) decreases to 0, showing that increasing *n* makes branching conditions more lenient. Oppositely, as *r* increases from 0 to 1, the rightmost term increases monotonically from 1*/n*, at which point branching conditions are maximally lenient, to 1, at which point the conditions cannot be met. Thus, increasing *r* makes branching conditions more stringent.

In the plane defined by *a*_1_ and *a*_2_, the total area *V* of points that satisfy (10) is a measure of how lenient or stringent the branching conditions are. In Figure 2 (left column), we plot how this area changes with *n* and *r*.

**Figure 1:**
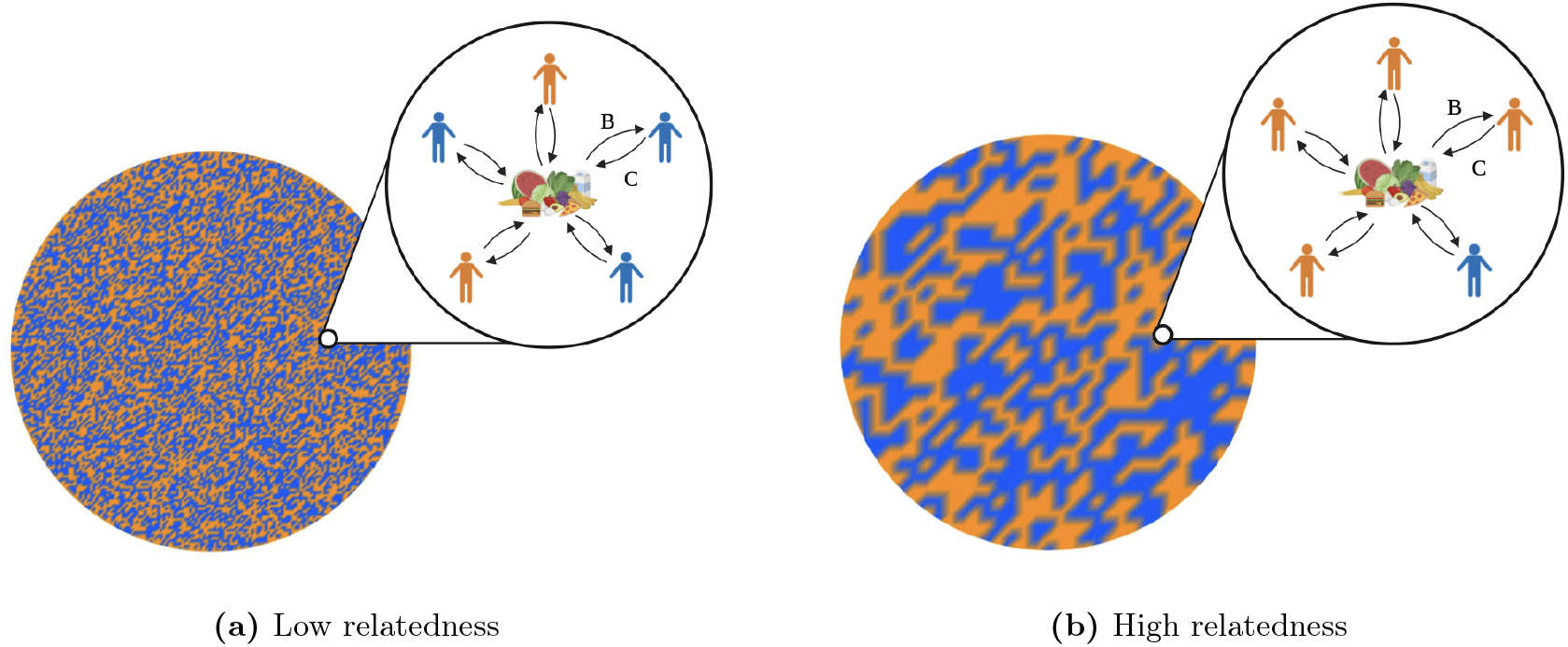
Public goods games with n = 5 in populations with different levels of relatedness. In this illustration, two phenotypes (blue and orange) exist in the populations, and individuals play the public goods game with their spatial neighbors. In the low-relatedness setting (a), the spatial distribution of types is approximately well-mixed, so the probability that a focal individual plays a game with either type is proportional to the frequency of that type in the overall population. In the high-relatedness setting (b), individuals of the same type have a higher probability of being near each other (proportional to the parameter *r*), so a focal individual is more likely to play the public goods game with coplayers who share its type.

**Figure 2:**
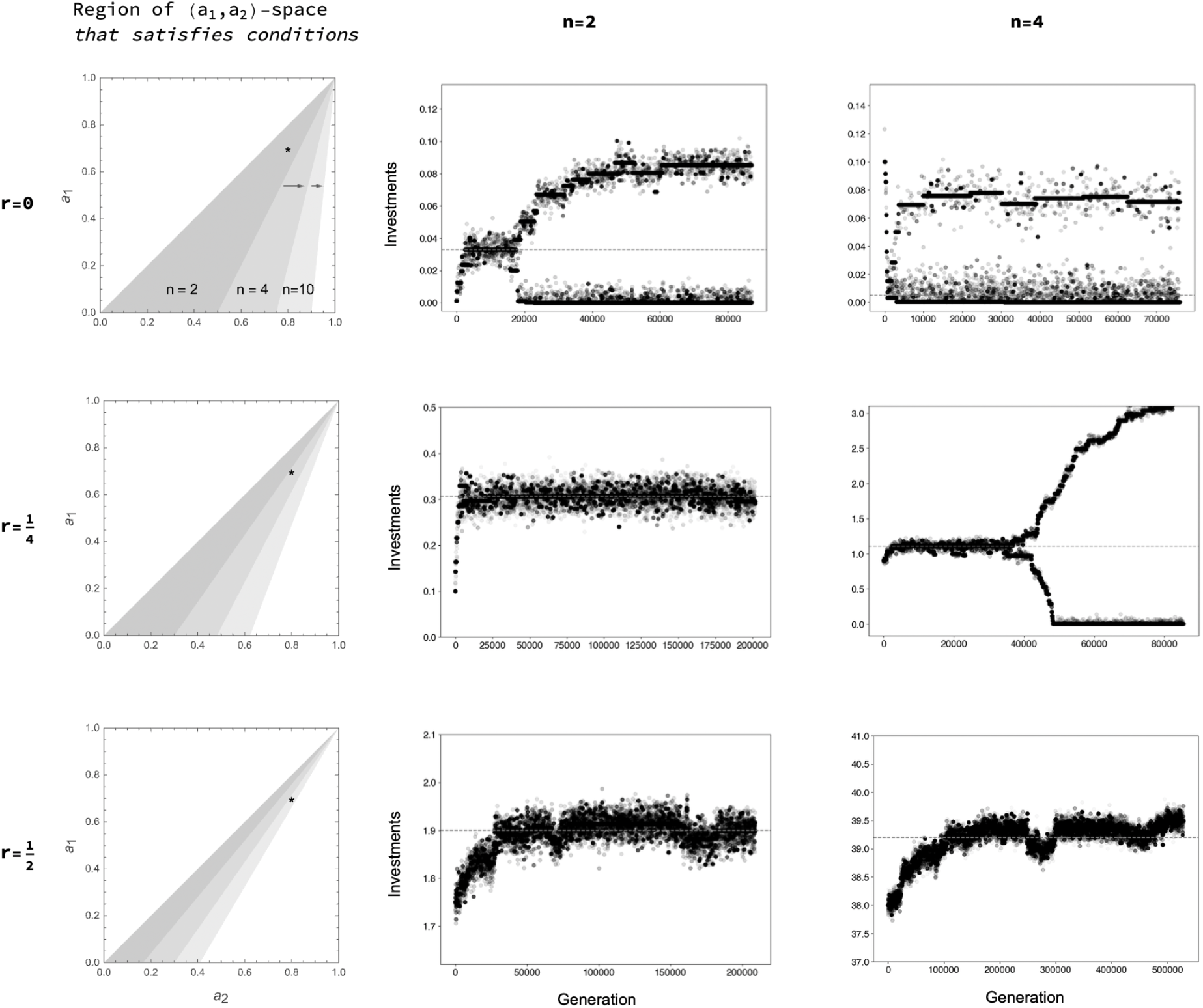
Evolutionary branching depends on group size *n* and relatedness *r*. The left column shows the area of (*a*_1_, *a*_2_)-space in which the parameters satisfy evolutionary branching conditions (9), as a function of *n* and *r*. The lighter-shaded areas indicate that higher levels of *n* (as labeled) are required to satisfy the conditions. Holding *r* constant, increasing *n* increases the area, whereas holding *n* constant, increasing *r* decreases it. The right two columns show sample simulations of investments in a public good for each labeled combination of *n* and *r*. We used a Moran process for the simulations, with fitness proportional to expected payoff. For all runs, *a*_1_ = 0.7 and *a*_2_ = 0.8. This parameter combination implies that investments will exhibit either evolutionary branching or continuous stability, depending on *n* and *r*, as indicated by the asterisk in the left column plots. The gray dashed lines indicate the monomorphic critical investment *z*_*_ for each parametrization.

The critical investment *z*_*_ for power function payoffs is

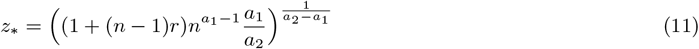

Note that 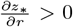, corresponding to our general result from section 3.3. The effect of *n* on *z*_*_ depends on two terms in (11): (1 + (*n* − 1)*r*), which shows that *n* scales the effect of assortment *r*; and 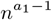, which is due to the implicit effect of *n* on public benefit *B*. With *r* > 0 and 0 < *a*_1_ < 1 (10), the former term increases with *n*, whereas the latter decreases with *n*. This leads to nonmonotonicity with small values of *r*, such that as *n* increases above 2, *z*_*_ decreases at first, reaching a minimum for some intermediate value of *n*, and then increases for higher *n* as the assortment term (1 + (*n* − 1)*r*) increases above the payoff term 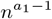. By contrast, for *r* = 0, *z*_*_ decreases monotonically with increasing *n* due to the payoff term; and for high values of *r* (∼ 0.5), *z*_*_ increases monotonically with *n*, because the assortment term always exceeds the payoff term.

#### 3.3.2 Quadratic Payoff Model

To check the robustness of these results, we carried out the same analysis assuming that public benefit and private cost are quadratic functions (equation (7); S.I. B.8.2). In this case, the concavities of the public benefit and private cost functions are constants. The evolutionary branching conditions under the quadratic payoff model simplify to

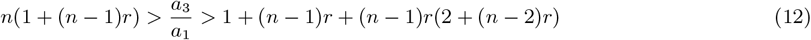

Unlike for power-function payoff, the volume of points that satisfy conditions (12) in the plane defined by concavity parameters *a*_1_ and *a*_3_ is unbounded when 0 ≤ *r* < 1. Additional parameters *a*_2_ and *a*_4_ must be chosen to compute the critical investment *z*_*_ and ensure that it is feasible. However, the range of values of the ratio *a*_3_*/a*_1_ that satisfy the branching conditions is delineated by the left- and right-hand terms in (12), which depend only on *n* and *r*. Taking the difference *D* between these terms, we obtain *D* = *n*(1 + (*n* − 1)*r*) − 1 + (*n* − 1)*r* + (*n* − 1)*r*(2 + (*n* − 2)*r*) = (*n* − 1)(1 +(*n* − 3)*r* − (*n* − 2)*r*^2^) = (*n* − 1)(1 − *r*^2^ +(*n* − 3)(*r* − *r* ^2^)). We can show that as *n* increases, *D* also increases, meaning that increasing *n* always makes branching conditions more lenient, as in the case of power-function payoffs. By contrast, the rate of change of *D* with respect to *r* (i.e.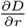) is nonmonotonic, switching from positive to negative at the value 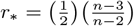. When *r* < *r*_*_, increasing *r* will also increase *D*, meaning that more assortment actually makes branching conditions more lenient, unlike for power-function payoffs. When *r* > *r*_*_, increasing *r* will decrease *D*, making branching conditions more stringent, as with power-function payoffs.

### 3.4 Intuition for the effects of group-size and assortment

To build intuition for the contrasting effects of *n* and *r* on the evolutionary branching conditions, consider a dimorphic population after branching, with two types, *u* and *v*, with investments *z*_*u*_ > *z*_*v*_, and frequencies *p* ∈ [0, 1] and 1−*p*. Typically, the dimorphic investments *z*_*u*_ and *z*_*v*_ are on opposite sides of the monomorphic critical investment *z*_*_. The selection gradient for a monomorphic population with resident investment *z*_*v*_ (*z*_*u*_) is positive (negative), due to the convergence stability condition (inequality (9), right). This implies that in the dimorphic population, the more frequently a mutant of type *z*_*v*_ (*z*_*u*_) plays the PG game with other *z*_*v*_ (*z*_*u*_) individuals, the more positive (negative) the selection gradient for *z*_*v*_ (*z*_*u*_) (S.I. C.1). This increase in stabilizing selection works against evolutionary branching, which requires the selection gradient for *z*_*v*_ (*z*_*u*_) to be negative (positive).

This reveals the intuition for why a sufficient increase in *r* makes evolutionary branching conditions more stringent: in a dimorphic population, increasing *r* makes the groups of individuals playing the PG game closer to a monomorphic group, from the perspective of either type.

It also suggests an inverse intuition for the effect of *n* (and of small *r* for quadratic payoff model): in a dimorphic population, increasing *n* reduces the variance of Σ*z*_*j*_*/n* around the mean trait value, reducing the probability density of *n*-groups that are monomorphic, from the perspective of either type. Thus, increasing *n* reduces the strength of stabilizing selection by making groups of *n* players less homogeneous on average. This makes it apparent that increasing *n* should make branching conditions less stringent. However, *n* does not only effect stabilizing selection.

Increasing the probability of expected *n*-groups, which are dimorphic, has the opposite effect on the selection gradient for either type as increasing the probability of homogeneous *n*-groups. This is due to the evolutionary instability condition, which requires that the monomorphic selection gradient has a minimum at *z*_*_. A subtlety of this condition is that a monomorphic population with a resident trait close to *z*_*_ is correspondingly close to a minimum of the selection gradient. Therefore, if the expected *n*-group in a dimorphic population is close to *z*_*_, then the lower and upper traits *z*_*v*_ and *z*_*u*_ are positioned on either side of this minimum, with their respective selection gradients pointing away from *z*_*_. Increasing *n* makes the expected *n*-group more likely, which strengthens the effect of this disruptive selection on *z*_*v*_ and *z*_*u*_. When the dimorphic traits are a small distance from *z*_*_ (and each other), some level of assortment can feasibly have this same effect, which explains the contrasting effect of small values of *r* with the quadratic payoff model. Overall, increasing *n* makes branching conditions more lenient.

### 3.5 Stability properties of dimorphic critical points with one public good

Next we turn to the long-term outcome after branching. The adaptive dynamics of the dimorphic population admits a critical point. As stated in section 2.1, this point (*z*_*u*_, *z*_*v*_)* may be both convergently and evolutionarily stable, in which case it is the final outcome of evolution, or it may be convergently stable but evolutionarily unstable, in which case one or both of the dimorphic types can be invaded by nearby mutants, resulting in further evolutionary branching into three or more types.

The dimorphic selection gradient with power-function payoffs has a critical point when evolutionary branching conditions are satisfied (i.e. when *a*_1_ and *a*_2_ are in the shaded region in the left column of Figure 2; S.I. B.9). We computed the dimorphic critical point (*z*_*u*_, *z*_*v*_)* numerically for a large sample of parametrizations and evaluated the stability properties of each point (Figure 3, S.I. B.9). All dimorphic critical points are convergently stable. With no relatedness (*r* = 0), they are also evolutionarily unstable. If *a*_2_ is only slightly greater than *a*_1_, both resident types can be invaded by nearby mutants, resulting in four existing types after a secondary branching; if *a*_2_ exceeds *a*_1_ by a greater amount, the resident type with the lower investment level can be invaded by nearby mutants, resulting in three types. With positive relatedness (*r* > 0), a subset of the critical points in (*a*_1_, *a*_2_)-space becomes evolutionarily stable, meaning they are the end-point of the adaptive dynamics (given *a*_1_ and *a*_2_). For this subset, *a*_2_ exceeds *a*_1_ by the maximum possible amount, while staying within the parameter region that allows branching.

**Figure 3:**
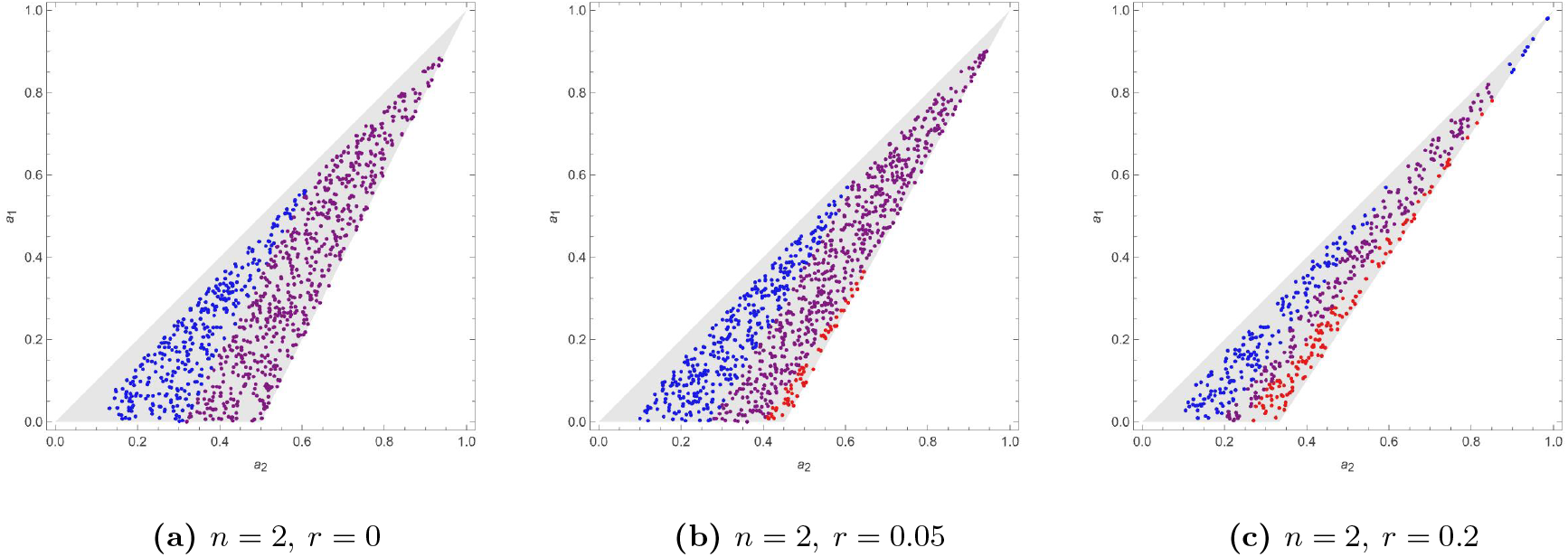
Stability properties of dimorphic critical points with power-function payoffs. Each dot in (*a*_1_, *a*_2_)-space corresponds to the dimorphic critical point given those parameters. Blue dots indicate that both dimorphic residents are evolutionarily unstable, leading to four types that emerge after secondary branching event(s). Purple dots indicate that only the dimorphic resident with the lower investment is evolutionarily unstable while the other resident is stable, implying three existing types after the secondary branching event, two of which contribute much less than the resident with the higher investment. Red dots indicate that both dimorphic residents are evolutionarily stable, implying that the dimorphism is the end-point of the adaptive dynamics. We only observed the latter case when *r* > 0. The gray-shaded area in either subfigure depicts the region of parameter space that meets evolutionary branching requirements, given group size *n* and relatedness *r*. For parametrizations with small values of *a*_2_ − *a*_1_ (left side of the branching region) we could not compute the dimorphic critical point, because the critical investment of the lower-investing type became so small that our numerical solver could not evaluate it accurately (S.I. B.9).

Thus, positive relatedness among individuals can stabilize division of labor at two types, and this happens in contexts where the public benefit diminishes much more slowly than the private cost (*a*_1_ << *a*_2_) at the critical point. Since increasing *r* makes the monomorphic critical point more evolutionarily stable, it is natural that it has a similar stabilizing effect on the dimorphic critical point. The same qualitative role of relatedness appears when payoffs are written as the sum of two public goods.

Evolution of investments after branching under the quadratic payoff model also has a dimorphic end-point, but for a different reason than for the power-function model. With the quadratic model, the selection gradient for *z*_*u*_ and *z*_*v*_ does not have a critical point, but the investment values have natural bounds (i.e. 0 and the lower of the maxima of *B* and *C*). After branching, the higher (lower) investment evolves to the upper (lower) boundary and is constrained to stability by the positive (negative) selection gradient. Further evolutionary stability analysis is not applicable in this case.

### 3.6 Evolutionary branching for two public goods

The simplest way to define fitness when there are two public goods is to add the payoffs of the two goods together. This leads to a payoff function of the form *π* = *B*_1_ − *C*_1_ + *B*_2_ − *C*_2_, where *B*_1_ (*B*_2_) and *C*_1_ (*C*_2_) are the public benefit and private cost functions for good 1 (good 2), respectively. A further simplifying assumption is that the public benefit and private cost of good *k* are functions only of investments in good *k*. In this sense, the two goods are substitutable, and there are no interactions between them. Let *z*_*i,k*_ denote an individual *i*’s investment in good *k*. The adaptive dynamics of investments in this setting with no assortment (*r* = 0) have been studied by Henriques et al. (2021) and Fielding et al. (2025).

Of particular interest is the case in which investments in both goods undergo evolutionary branching — that is, when (*B*_1_, *C*_1_) and (*B*_2_, *C*_2_) each satisfy the single-good branching conditions (9). After branching, the population consists of two types, *u* and *v*, each characterized by a pair of investments. The subsequent adaptive dynamics can lead to two qualitatively distinct outcomes. In specialization, one type invests more in good 1 and less in good 2, and the other does the opposite. In exploitation, one type invests above the critical level in both goods, while the other invests below the critical level in both. These outcomes are bistable: which one arises depends on the initial configuration of the dimorphism that forms at the branching point. The mathematical reason is straightforward. Once the direction of branching is fixed for good 1 (for example, type *u* invests high and type *v* low), there remain two possible orientations for branching on good 2 (S.I. C.1). These alternative orientations correspond precisely to exploitation versus specialization.

Importantly, the qualitative results described above hold under partial assortment (0 < *r* < 1). Perhaps more interesting is that dimorphic critical points 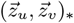exist and are evolutionarily stable with positive assortment (*r* > 0) for power-function payoffs (S.I. C.2). These, finally, are continuously stable divisions of labor that emerge from a homogeneous population. As discussed in Section 3.4, the intuition is that assortment primarily alters the stability of the critical point, without changing the qualitative pattern of branching itself. The two investments still diverge after branching; increasing *r* simply determines whether that divergence continues or settles at a stable dimorphism.

## 4 Discussion

In this article, we extended the theory of division of labor due to evolutionary branching of investments in public goods by incorporating assortment of related individuals. The conditions for branching (9) are functions of public-good group size *n*, relatedness *r*, and the concavities of the public benefit *B* and private cost *C* functions with respect to all of their investment inputs. Overall, we found that increasing group size makes these conditions more lenient, while increasing relatedness tends to make them less lenient but may have the opposite effect depending on how payoffs are defined. We also showed that public benefits must diminish with respect to all investments for branching to occur, for a preponderance of reasonably-defined benefit functions; moreover, if payoffs are defined as the difference between public benefit and private cost (rather than their product), then the cost function must diminish more quickly than the benefit function with respect to investment for branching to occur.

Our results suggest candidate ecological settings in which biologists should expect to observe organic division of labor. Populations in which public goods are shared within large groups of individuals with low relatedness are more likely to exhibit evolutionary branching. This applies to many bacterial populations, which share public goods with large numbers of neighbors via diffusion and may have limited assortment of related individuals due to motility [Cordero et al., 2012]. It also applies to ancestral humans, who often shared public goods with large groups and had a relatively low probability of interacting with kin [Apicella and Silk, 2019].

Division of labor is also more likely in natural populations when the benefits and costs derived from a public good both diminish with respect to their investment inputs, and especially when costs diminish faster than benefits. The latter prediction holds when public benefits are a function of the sum or product of investments (*etc*. S.I. B.4) and payoff is equal to the difference between benefits and costs.

Saturating functions are arguably more realistic than linear or accelerating ones for the fitness benefits of public goods in natural populations because at high public good levels, fitness benefits are likely to be limited by outside factors. In the classic siderophore example, a bacterium’s rate of iron uptake diminishes as a function of the concentration of secreted siderophores in its extracellular environment, as more iron is extracted [Bergeron and Weimar, 1990]. Similarly among human and other animal foragers, the rate at which food is acquired diminishes as a function of group members’ collective search effort, as the concentration of food in the environment is depleted [Eccard and Leisenjohann, 2008]. Saturating costs, in turn, can arise when phenotypic plasticity or learning allows individuals to reduce the marginal cost of their investments when investing at high levels. For example, it is possible for bacteria to plastically increase the efficiency of the cellular machinery used to produce metabolites, reducing their marginal cost [Mullen et al., 2015]. Similar marginal cost-reducing effects can arise in human foraging, as individuals learn to improve their search and handling techniques. Thus, empirical measurement of the relative concavities of public benefits and private costs can inform predictions about the likelihood of division of labor in natural populations.

Increasing assortment tends to decrease the strength of disruptive selection, making division of labor less likely. Evolutionary branching is possible with partial (0 < *r* < 1) but becomes impossible with complete assortment (*r* = 1). Partial assortment also makes it possible for the critical division of labor formed by the two types after branching to be evolutionarily stable, so not prone to further branching, whereas with no assortment the critical dimorphism is always unstable. Thus, assortment can stabilize the evolution of polymorphisms as well as homogeneous populations.

Our analysis builds on the evolutionary-branching literature, especially work on continuous investments in public goods. Doebeli et al. [2004] analyzed a single public good in the absence of relatedness, and subsequent studies extended the same framework to two substitutable goods in well-mixed populations [Fielding et al., 2025; Henriques et al., 2021]. To incorporate relatedness, we draw on a complementary tradition in quantitative genetics. Wakano and Lehmann [2014] derived branching conditions for general fitness functions under relatedness. Our condition (9) is a specialization of their framework to payoffs of the form “expected public benefit minus private cost,” with pairwise relatedness represented by a constant probability *r*. In Appendix S.I. B we re-derive (9) using standard adaptive-dynamics calculations, which makes explicit the equivalence between the adaptive-dynamics and quantitative-genetic routes to branching for quantitative traits. Lehmann and Mullon [2025] also provided a general treatment of evolutionary branching under relatedness. Their work demonstrates that the monomorphic critical point satisfies a marginal form of Hamilton’s rule [Hamilton, 1964].

The division of labor on public goods we study here is distinct from reproductive division of labor. In many models of reproductive division of labor, there is a (potentially stochastic) assignment of individuals (e.g., individual cells in a colony) into classes that then might evolve class-dependent strategies [Cooper et al., 2022; Michod, 2006, 2007; Rueffler et al., 2012]. Instead, our evolutionary branching model starts with all individuals being symmetric, without differences in class. The differences in investment strategies evolve through the shape of the tradeoffs determining the costs and benefits of the investments. A related distinction is that our group formation process assigns individuals randomly (possibly with assortment) into groups whereas in the class-structured groups of reproductive division of labor literature, a mechanism usually constrains the group composition. These qualitative distinctions between the ecological and reproductive theories of division of labor correspond to our result that evolutionary branching becomes impossible with complete relatedness. In that setting, a class-assignment mechanism is required for labor division, as in the reproductive theory.

Our study has several limitations that point to future directions. Most importantly, the adaptive-dynamics framework assumes that mutations are infinitesimal: a mutant differs only marginally from the resident. This assumption is appropriate when traits are polygenic, so that small genetic changes produce small phenotypic shifts [Waxman and Gavrilets, 2005], or when traits change through behavioral imitation with small copying errors [Nowak and Sigmund, 2004]. Many biological and cultural traits plausibly fit this description. For example, microbial production of siderophores depends on the coordinated expression of many genes [Crosa, 1989], and imitation shapes behavior in humans and other animals [Byrne and Resson, 1998; Gariepy et al., 2014; Hanna and Meltzoff, 1993]. However, not all traits change gradually. Some public-good investments may depend on one or a few loci, in which case mutations can produce large phenotypic jumps. In such settings, investment is better modeled as a discrete or binary trait (e.g., producer versus non-producer), rather than as a continuously varying quantitative trait. Extending the present analysis to incorporate large-effect mutations or discrete trait spaces would clarify how robust our branching results are beyond the small-mutation limit.

Our study also makes several simplifying assumptions to isolate the mechanisms of interest. First, we treat public-good group size *n* as fixed. In natural populations, group size is typically variable and better described by a distribution [James, 1953; Peña, 2011]. Allowing group size to fluctuate would add another layer of ecological heterogeneity that could interact with the branching conditions derived here. Similarly, we treat relatedness *r* as an exogenous parameter. In reality, relatedness can evolve through demographic processes such as dispersal or spatial structure [Wakano and Lehmann, 2014]. By holding *r* fixed, we isolate its direct effect on the curvature conditions that determine evolutionary branching, rather than conflating those effects with the ecological processes that generate relatedness.

We also assumed that the benefits of the two public goods do not interact. In our model, the public benefit of one good depends only on investments in that good, and not on investments in the other. This provides a clean baseline against which more complex interactions can later be evaluated. In many biological systems, however, public goods are complementary. In mound-building mice, for example, transporting vegetation and transporting roof-building material contribute to different components of the same structure [Hurtado et al., 2013]. The effectiveness of the mound depends on both tasks, so the benefit of one is implicitly tied to investment in the other. Likewise, in human hunter–gatherer societies, meat and plant foods provide complementary nutritional resources [Armelagos, 2014]. In such cases, overall fitness gains depend jointly on investments across goods. Incorporating such interactions—whether synergistic or antagonistic—would be a natural extension of the present framework. Doing so would allow division of labor to arise not only from curvature in costs and benefits, but also from complementarities between tasks.

## Supporting information

Simulation output

## A Adaptive dynamics of investments in one public good

We derive the conditions for convergence stability and evolutionary instability of a monomorphic critical investment under a simple relatedness scheme. Our basic assumption is that fitness for a focal individual is given by the expected payoff from a public goods game with *n* − 1 other players, each of whom is the same type as the focal individual with probability *r* ∈ [0, 1] (or is a randomly drawn type with probability 1 − *r*).

### A.1 Interchangeable benefit functions

Define public benefit as a vector-valued function of *n* investment traits, that is 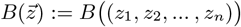.

#### Definition 1

For any *i, j* ∈ (1, …, *n*) s.t. *i* ≠ *j*, let *z*_i_ = *x* and *z*_*j*_= *y* where *x, y* ∈ ℝ. Investments are *interchangeable* iff. 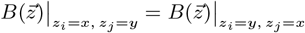

We can show that definition 1 implies the following equalities:

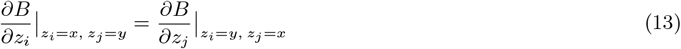

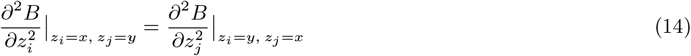

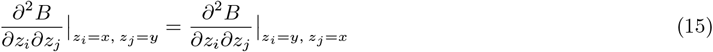

That is, if the investments of two individuals among the *n* switch, the first and second partial derivatives of *B* with respect to their investments also switch. We use the definition of interchangeability to define fitness (16). We use the first interchangeability property ((13)) above to derive the selection gradient (18). And, we use the latter properties ((14)) and ((15)) above to derive the convergence stability condition (19) and the evolutionary instability condition (22).

### A.2 Fitness

The fitness *w* of a mutant with trait *z*_mut_ in a population with resident trait *z*_res_ is

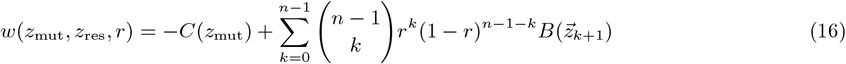

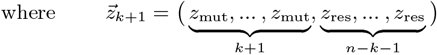

That is, 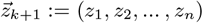 st. *z*_*i*_ = *z*_mut_ ∀*i* ≤ *k* + 1 and *z*_*i*_ = *z*_res_ ∀*i* > *k* + 1.

Note that

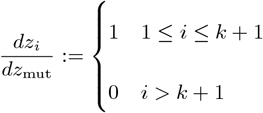

And 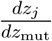 is defined analogously.

## B Evolutionary branching conditions

### B.1 Convergence stability

We derive the selection gradient as follows:

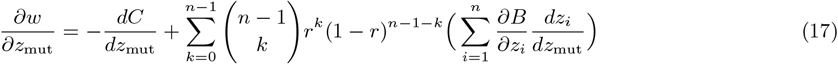

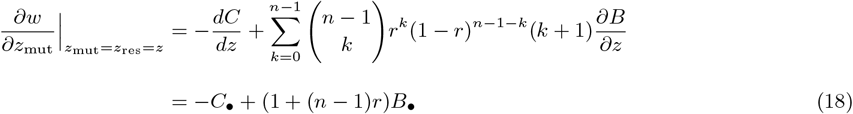

Where 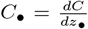 and 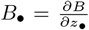 are the derivatives of *C* and *B* wrt. a focal investment (•) among the *n*, when all are evaluated at the resident point *z*. Note that the derivation of mutant fitness (18) assumes interchageability of investments as defined above.

The CS condition is

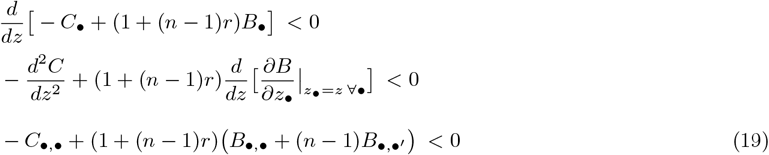

Where 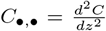is the second derivative of the cost function, 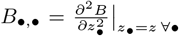is the second derivative of the benefit function wrt. any focal trait, and 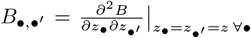is the cross-derivative of the benefit function wrt. focal trait *z*_•_ and any nonfocal trait *z*_•_′, where all are evaluated at resident trait *z*. Note that (19) is equivalent to inequality (14) in Wakano and Lehmann [2014].

### B.2 Evolutionary instability

Return to equation (17) above and take the second partial derivative of fitness wrt. *z*_mut_:

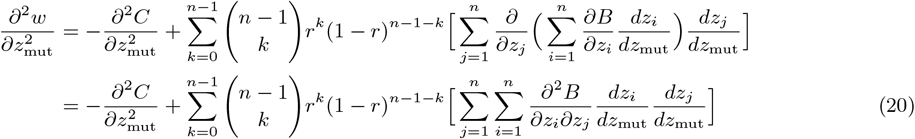

Now evaluate at a resident trait, such that *z*_mut_ = *z*_res_ = *z*:

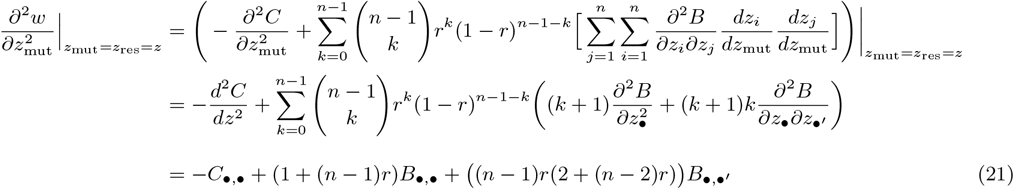

Where *z*_•_ and *z*_•_′ are generic focal and nonfocal traits, as defined above, and we evaluated the sum in Mathematica. After simplifying, we obtain the following evolutionary instability condition:

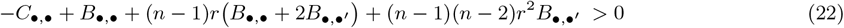

This is equivalent to inequality (29) in Wakano and Lehmann [2014], with 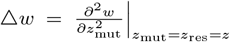, *w*_SS_ = *B*_•,•_ − *C*_•,•_, *w*_*SD*_ = 2(*n* − 1)*B*_•,•_′, *w*_*DD*_ = (*n* − 1)*B*_•,•_, *w*_*DD*_′ = (*n* − 1)(*n* − 2)*B*_•,•_′, *R*_2_ = *r, R*_3_ = *r*, and *△ r* = 0 by assumption.

### B.3 Branching requires that the cross derivative is negative (*B*_•,•_′ < 0) and relatedness is nonclonal (*r* < 1)

From CS condition (19) above, we have

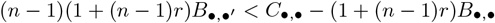

And from E(in)S condition (22) above, we have

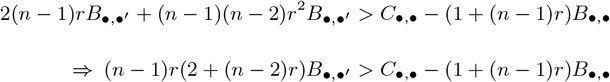

Combining these, we obtain

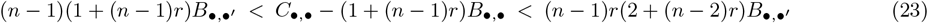

When *r* = 0, we obtain the standard evolutionary branching conditions with no assortment, which imply that costs, focal benefits, and cross-benefits must all be saturating, that is *C*_•,•_ < 0, *B*_•,•_ < 0, *B*_•,•_′ < 0.

When *r* ∈ (0, 1), we can easily show that (*n* − 1)(1 + (*n* − 1)*r*) *>* (*n* − 1)*r*(2 + (*n* − 2)*r*), implying that the cross-derivative must again be negative (*B*_•,•_′< 0) for evolutionary branching conditions to be met (from the left and right sides of (23)). When *r* = 1, we obtain *n*(*n* − 1)*B*_•,•_′ < *C*_•,•_ − *B*_•,•_ < *n*(*n* − 1)*B*_•,•_′ which is false, so evolutionary branching cannot happen with *r* = 1.

### B.4 General class of public benefit functions for which branching implies diminishing *B, C*

Our basic inference from the general branching conditions (23) is that *B*_•,•_′ < 0. For additive or combinatorial public benefit functions (defined below), we can further show that *B*_•,•_ *≤ B*_•,•_′, which in turn implies that *B*_•,•_, *C*_•,•_ < 0. In this section, we define the generic properties of *B* that allow us to make this inference.

Let *B* : 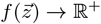, where we assume 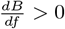 and *∀j* ∈ (1, …, *n*),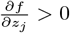, i.e. the effect on *B* of each individual’s investment is positive. We have

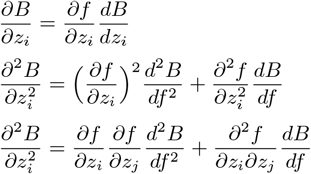

The latter two terms, evaluated at the monomorphic critical point, are *B*_•,•_ and *B*_•,•_′.

Note that 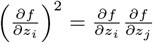 due to interchangeability of investments, and 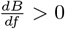 by assumption.

Therefore, if 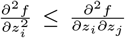, then *B*_•,•_ *≤ B*_•,•_′. The standard additive function 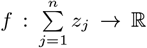 satisfies this requirement. More generally, any function *f* that is a sum of products of all combinations of *k* investments, or a sum of these sums for multiple *k* values, satisfies this requirement (where *k* ∈ {1, …, *n*}). If *k* = 1, then *f* is the canonical function above; if *k* = *n*, then *f* is the product of all *n* investments; and for example, if *k* = 2, then *f* is the sum of all pairs of investments, that is 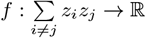.

We can construct a counterexample *f* function for which investments are interchangeable and yet 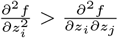 : Let 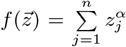. Then for any *i* and *j* from 1 to *n* with *i ≠ j*, we have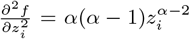,and 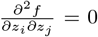 Therefore if *α >* 1, then 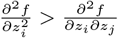and evolutionary branching does not imply *B*_•,•_′, *B*_•,•_, *C*_•,•_ < 0. If 0 *≤ α ≤* 1, then 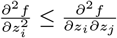, so evolutionary branching implies *B*_•,•_′, *B*_•,•_, *C*_•,•_ < 0.

### B.5 Other observations

By contrast, if the monomorphic critical point is convergently unstable and evolutionarily unstable, we cannot infer the sign of *B*_•,•_′ or the signs of *B*_•,•_ of *C*_•,•_. This is because in the unstable setting, the conditions for the second partial derivatives can be found by moving *both* the LHS and RHS of branching conditions (23) to the right (left) of the middle term. Under these different conditions, there is no information about the order of the LHS and RHS, which under evolutionary branching conditions (23) implies that *B*_•,•_′ < 0 and therefore *B*_•,•_, *C*_•,•_ < 0.

The same analysis applies for multiplicative payoff functions of the form *π*_i_ = *B · C* (see S.I. D below). We can show that if public benefits are in the additive/combinatorial family of functions, then *B*_•,•_′, *B*_•,•_ < 0 are necessary for evolutionary branching to occur.

### B.6 Effect of assortment *r* on critical point *z*_*\**_

The conditions for evolutionary branching imply that 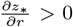, that is, higher levels of assortment increase the critical level.

Proof: The monomorphic critical point *z*_*\**_ can be found by solving

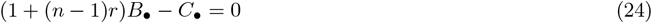

for resident trait level *z*. Write *z*_*\**_ := *z*_*\**_ (*n, r*) as a function of *n* and *r*, and at that point, write *B*_•_ := *B*_•_ (*z*_*\**_, …, *z*_*\**_) and *C*_•_ := *C*_•_ (*z*_*\**_) as functions of *z*_*\**_. Now take the total derivative of (24) with respect to *r*:

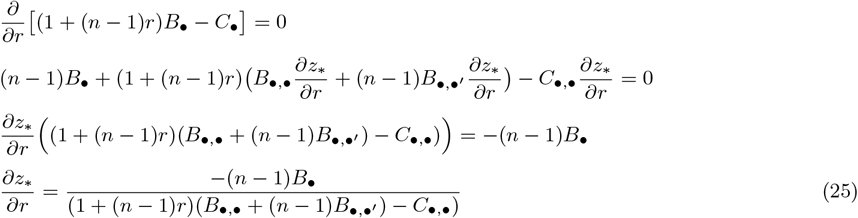

The numerator in (25) is negative because we assume *B*_•_ *>* 0, and the denominator is negative under the general branching conditions (23). This implies that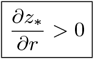.

### B.7 Branching conditions when *B* is a function of additive investments

If *B* is a function of the sum of investments Σ *z*_*j*_, then *B*_•,•_ = *B*_•,•_′. In this case, we have

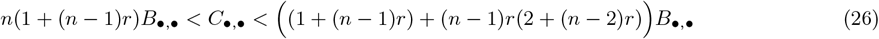

Because 1 + (*n* − 1)*r >* 2*r* + (*n* − 2)*r*^2^, the RHS of (26) is greater than the LHS *iff. B*_•,•_ < 0, which further implies that *C*_•,•_ < 0 in this case. We can divide by *B*_•,•_ to obtain

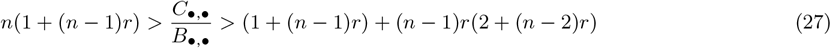

This pair of inequalities is used in the following section.

### B.8 Analysis with numerical payoff functions

#### B.8.1 Model 1

Let 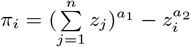. We have

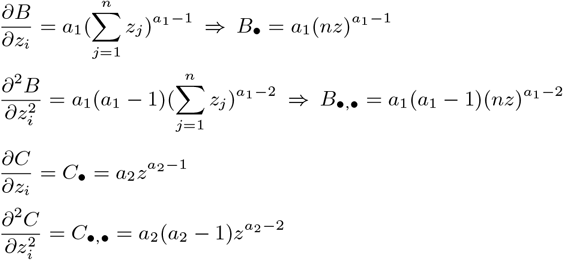

We solve for the critical point *z*_*\**_ by setting (1 + (*n* − 1)*r*)*B*_•_ − *C*_•_ = 0 and solving for *z*, to obtain

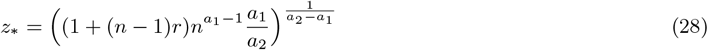

Note that

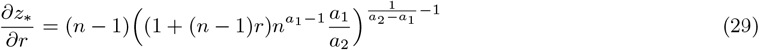

Equation (29) and our baseline assumptions that *a*_1_, *a*_2_ *>* 0 imply that 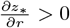.

Plug the expressions for *B*_•,•_ and *C*_•,•_ into (27) to obtain evolutionary branching conditions for model 1:

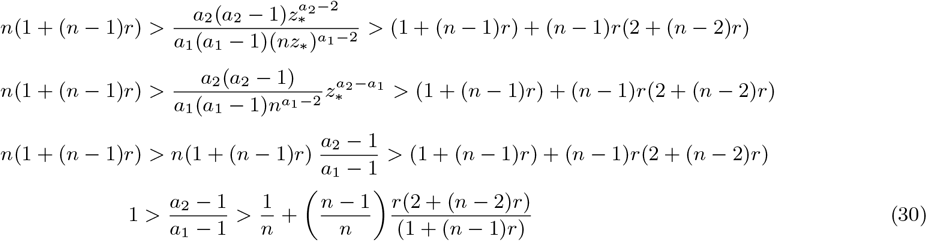

As *r* increases, the region of (*a*_1_, *a*_2_)-space that allows evolutionary branching becomes smaller. The area of this region is directly proportional to the quantity

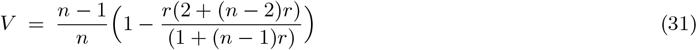

We can show that *V* increases with *n* by evaluating the quantity

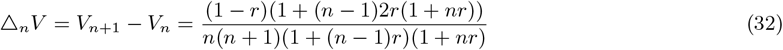

Each term in this quotient is positive, so *△*_n_*V >* 0.

The rate of change of *V* with respect to *r* is

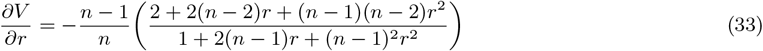

The term in parentheses is strictly positive, so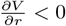.

Finally, we can examine the cross-effects of *n* and *r* by computing the following

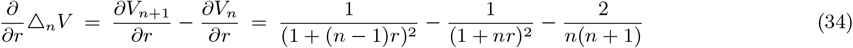

This expression is typically negative, meaning that for a given *r*, if *n* increases then the effect on *V* of increasing *r* will become more negative. However, for *n ≥* 6, there is a range of small *r* values for which the expression is positive, meaning that increasing *n* dampens the negative effect of *r* on *V*.

#### B.8.2 Model 2

Let 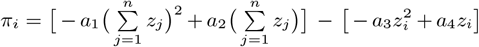. The first term in square brackets is public benefit, and the second (subtracted) bracketed term is private cost. We have

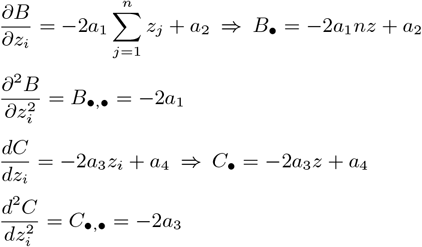

From equation (27), the evolutionary branching conditions for model 2 are as follows:

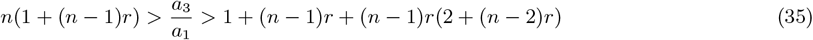

The region of (*a*_1_, *a*_3_)-space that allows evolutionary branching is a triangle in the upper-right quadrant that is defined by the LHS and RHS of (35). The area of the region is directly proportional to the difference *D* between these two quantities:

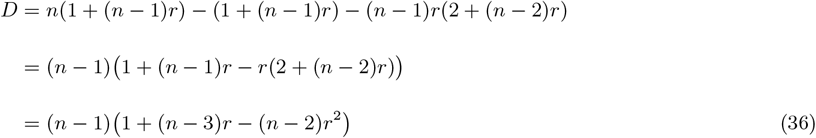

Show that *D* increases with *n* (i.e. *△*_n_*D >* 0):

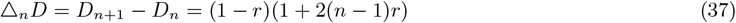

And the rate of change of *V* wrt. *r* is

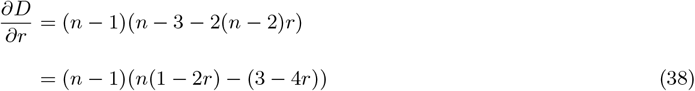

With *n* = 2, 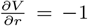with *n* = 3, 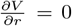 for *r* = 0 and decreases linearly with *r*; and with *n ≥* 4, 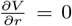 at 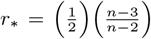 and decreases linearly with *r*, meaning that 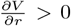 for *r* < *r*_*\**_ and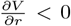 for *r > r*_*\**_. The cross-effects of *n* and *r* are given by

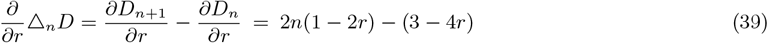

Which are linearly decreasing with *r*, similar to 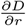.

### B.9 Adaptive dynamics with two types, with one public good and assortment (*r >* 0)

After evolutionary branching, there are two resident types, *u* and *v*, with traits *z*_*u*_ (*z*_*v*_) and frequencies *p* (1 − *p*), where *p* ∈ (0, 1).

Consider a mutant with investment *z*_mut_ in a resident population with investments (*z*_*u*_, *z*_*v*_). We can write the payoff to the mutant from the public goods game as

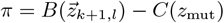

where 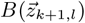 is a vector of which *k* + 1 elements are equal to *z*_mut_, while *l* elements are equal to *z*_*u*_ and *n* − 1 − *k* − *l* elements are equal to *z*_*v*_.

The fitness *w* of a mutant with trait *z*_mut_ in a population with resident traits (*z*_*u*_, *z*_*v*_) is

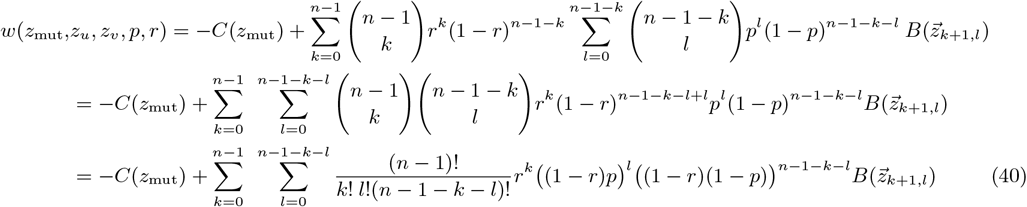

Thus, the public benefit 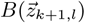 that the mutant receives from the game has a trinomial distribution. For each nonfocal player *j*, with probability *r, j* has trait *z*_mut_; with probability (1 − *r*)*p*, has trait *z*_*u*_; and with probability (1 − *r*)(1 − *p*), has trait *z*_*v*_.

The fitness of a type *u* individual in the resident population is written *w*(*z*_*u*_, *z*_*u*_, *z*_*v*_, *p, r*), i.e. treating the focal ‘mutant’ as a type *u* individual in (40) (fitness of type *v* is defined analogously).

The dynamics of *p* (the frequency of type *u*) are given by

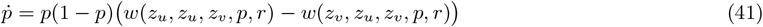

For the summed-investment family of payoff functions, if an internal equilibrium frequency *p*_*\**_ ∈ (0, 1) satisfying 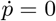 exists, then it is dynamically stable and unique. We prove this below in S.I. F. This allows us to initialize dimorphic investments *z*_*u*_ and *z*_*v*_ close to a branching point, where they have a unique frequency equilibrium *p*_*\**_ (*z*_*u*_, *z*_*v*_), and derive their adaptive dynamics by integrating the two-type selection gradient along the manifold defined by *p*_*\**_.

Note: if we distinguished relatedness within resident types (*r*_type_) from that within focal strains (*r*_mut_), we could compute the resulting adaptive dynamics by substituting *r*_type_ into (41) in our solution for *p*_*\**_, and substituting *r*_mut_ in our equation for mutant fitness (40).

The two-type selection gradient is computed by taking the partial derivative of fitness with respect to the mutant trait and evaluating at *z*_mut_ = *z*_*u*_ (for type *u*) and *z*_mut_ = *z*_*v*_ (for type *v*) when the resident traits *z*_*u*_ (*z*_*v*_) are at equilibrium frequencies *p*_*\**_ (1 − *p*_*\**_):

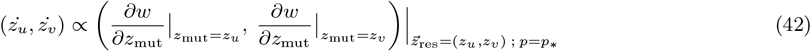

We derive an adaptive dynamical trajectory of *z*_*u*_ and *z*_*v*_ by choosing initial traits (*z*_u0_, *z*_v0_) with internal frequency equilibrium *p*_*\**0_ and integrating the two-type selection gradient (42). In every such case, if *z*_u0_ *> z*_v0_, then *z*_*u*_ evolves upward and *z*_*v*_ downward, and vice versa. The resident traits may evolve to a dimorphic critical point, at which 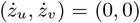, or to a point at which *p*_*\**_ = 0 or 1, so one type fixes and the population reverts to a monomorphic state.

Finally, to analyze the convergence and evolutionary (in)stability of a dimorphic critical point, we define the Hessian and Jacobian matrices of the selection gradient, following Leimar [2009]:

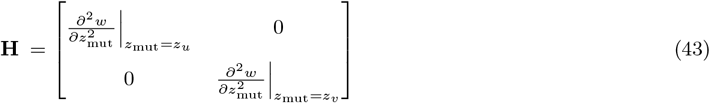

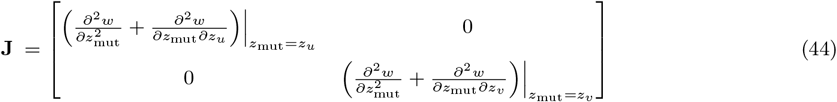

Where all elements of both matrices are evaluated at the resident dimorphic point (*z*_*u*_, *z*_*v*_, *p*_*\**_).

If the matrix J is negative definite, then the dimorphism is convergence stable. If the matrix H is negative definite, it is evolutionarily stable. If H is positive definite or indefinite, further evolutionary branching into additional types will occur. It is helpful that the definiteness of either H or J depends entirely on the signs of the diagonal elements, which are their eigenvalues.

For a given specification of payoff *π*, such as the power-function or quadratic forms analyzed above, we can check if the monomorphic critical point satisfies branching conditions, solve for a dimorphic critical point (*z*_*u*_, *z*_*v*_)_*\**_ (if it exists) by setting (42) equal to 0, and determine the evolutionary and convergence (in)stability of (*z*_*u*_, *z*_*v*_)_*\**_ by computing (43) and (44), respectively.

We depict the evolutionary stability properties of a large sample of dimorphic critical points in figure 3 (main text) for the power-function form of payoff. For very small values of *a*_2_ − *a*_1_, such that the rate at which cost diminishes only slightly exceeds the rate at which benefit diminishes, the critical investment for the lower-investing type decreases to an extremely small value on the order of 10^−5^. This causes the magnitudes of the components of the selection gradient for both types to increase rapidly. This in turn makes numerical integration of the dimorphic selection gradient challenging, because integrating that of the lower type for even a very small finite step size can cause the lower trait to drop below 0, which is undefined and breaks the solver that we used. That is the reason for the blank areas in the left part of the branching region in figure 3. Better algorithms could be used to solve for more critical points in this left region.

## C Adaptive dynamics with two public goods

With two public goods, each individual now has two investment traits, such that for individual *i* ∈ (1, …, *n*), their investments are written 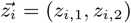.

The simplest way to define the payoff from two public goods is as follows. Let *B*_*k*_ and *C*_*k*_ denote the benefit and cost functions for good *k*, and let {*z*_−*i,k*_} denote the *n* − 1 nonfocal investments in good *k*. Payoff for *i* is

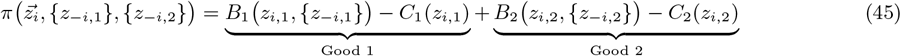

We label the two types that arise after branching *u* and *v*, with investment traits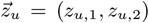 and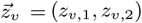and frequencies *p*, (1 − *p*). As above, the fitness of a mutant with trait vector 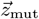 in this population is

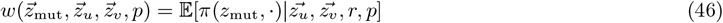

The selection gradient 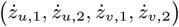 for each of the four investment traits is proportional to

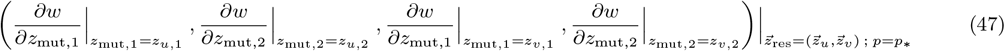

Where *p* = *p*_*\**_ is the solution of *w* 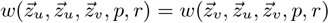 (see (41)).

If conditions (23) are true for either good 1 (*B*_1_, *C*_1_) or good 2 (*B*_2_, *C*_2_), then evolutionary branching into two types will occur. For payoff model 1, if (23) holds for both goods, the two types evolve to one of two dimorphic critical points. These points represent two characteristically different divisions of labor: exploitation (ex), in which some individuals invest more in both goods while others invest less in both; and specialization (sp), in which some individuals invest more in good 1 and less in good 2, while others invest more in good 2 and less in good 1. The exploitation and specialization outcomes are multistable. In our numerical investigations, the following basins for these outcomes hold for all parameters satisfying (23) (up to symmetry with *v*):

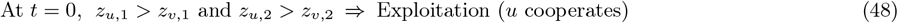

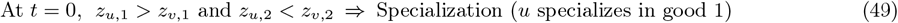

However, we have not analytically proved these basin boundaries.

### C.1 Initial dimorphisms after branching determine the division of labor over time

Adaptive dynamics describes the expected trait substitution sequence with infinitesimal mutations, which is a deterministic trajectory. As such, the canonical description of evolutionary branching is a monomorphic population bthat evolves precisely to the critical point *z*_*\**_, at which mutants on either side can invade and coexist with the resident, forming an initial dimorphism. At that point, the adaptive dynamics of the two types must be derived to assess further evolution (see above).

If the fitness function is sufficiently smooth, the two resident investments after evolutionary branching evolve away from each other. In this setting, a candidate initial dimorphism (*z*_*u*_, *z*_*v*_)t=0 may lie on one of these two-type trajectories. This becomes more likely as *z*_*u*_ (*t* = 0) and *z*_*v*_ (*t* = 0) get closer to the monomorphic critical point *z*_*\**_. However, for payoff model 1, *z*_*u*_ (*t* = 0) and *z*_*v*_ (*t* = 0) can *both* be above (or below) the critical point *z*_*\**_ yet lie on a two-type trajectory, where the trait that is closer to *z*_*\**_ evolves towards it.

In general, if a candidate initial dimorphism (*z*_*u*_, *z*_*v*_)t=0 is ecologically stable (i.e. *p*_*\**_ ∈ (0, 1)), then it lies on a dimorphic post-branching trajectory, and the lower (higher) investment will evolve downward (upward) for some amount of time. The preceding logic applies to the adaptive dynamics of investments in *two* public goods after evolutionary branching (see equations (45), (46) and (47)).

In turn, we can conclude that the division of labor (i.e. exploitation (35) or specialization (36)) that evolves after evolutionary branching on two public goods matches that of the initial dimorphism that appears: a population whose initial dimorphism has an exploitation DOL (*z*_*u*,1_ *> z*_*v*,1_ and *z*_*u*,2_ *> z*_*v*,2_ or vice versa) is characterized by exploitation at *t >* 0; a population whose initial dimorphism has a specialization DOL (*z*_*u*,1_ *> z*_v,1_ and *z*_*u*,2_ < *z*_v,2_ or vice versa) is characterized by specialization at *t >* 0.

### C.2 Defining the Hessian and Jacobian matrices for two goods and two types

Finally, following Leimar [2009] as before (see section B.9), we define the Hessian and Jacobian matrices of the selection gradient for two goods and two types.

The Hessian matrix is block diagonal with entries equal to the second partial derivatives of fitness with respect to both mutant traits, evaluated at the resident point. Following equation (7) from [Leimar, 2009], we can denote one of these blocks for type *k* ∈ {*u, v*} as follows:

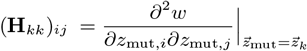

Where the first (second) partial derivative is taken with respect to a mutant of type *k*’s investment in public good *i* (*j*), with *i, j* ∈ {1, 2}. Note that because in our payoff function (45) investments in different goods affect different summands (i.e. there are no interactions between goods), we know that (**H**_*kk*_)_*ij*_ = 0 for *i ≠ j*.

This gives us the following Hessian matrix for two goods and two types (using column ordering (*z*_*u*,1_, *z*_*u*,2_, *z*_v,1_, *z*_v,2_)):

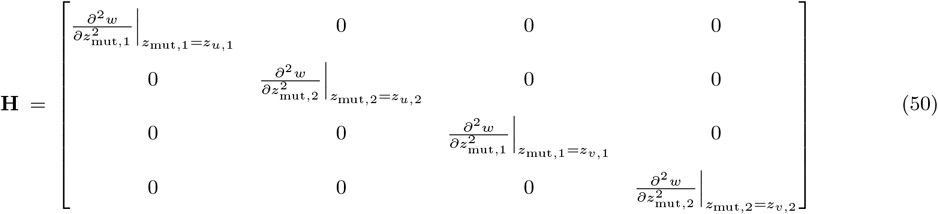

Where, as before with one good, all elements of *H* are evaluated at the resident dimorphic point 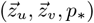. Again, the eigenvalues are the diagonal elements, and they determine the definteness of **H**. The same is not true for **J** below.

To define the Jacobian, we refer to equation (10) in [Leimar, 2009], which defines the matrix **Q** of cross-derivatives of fitness with respect to all mutant traits and all resident traits. We can denote a block of such cross derivatives where the mutant is from type *k* and the resident is from type *l* as follows:

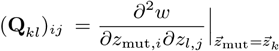

The desired Jacobian matrix is evaluated **J** = **H+Q**. We write this 4 *×* 4 matrix below in two blocks: the left half (4 *×* 2) and the right half (4 *×* 2).

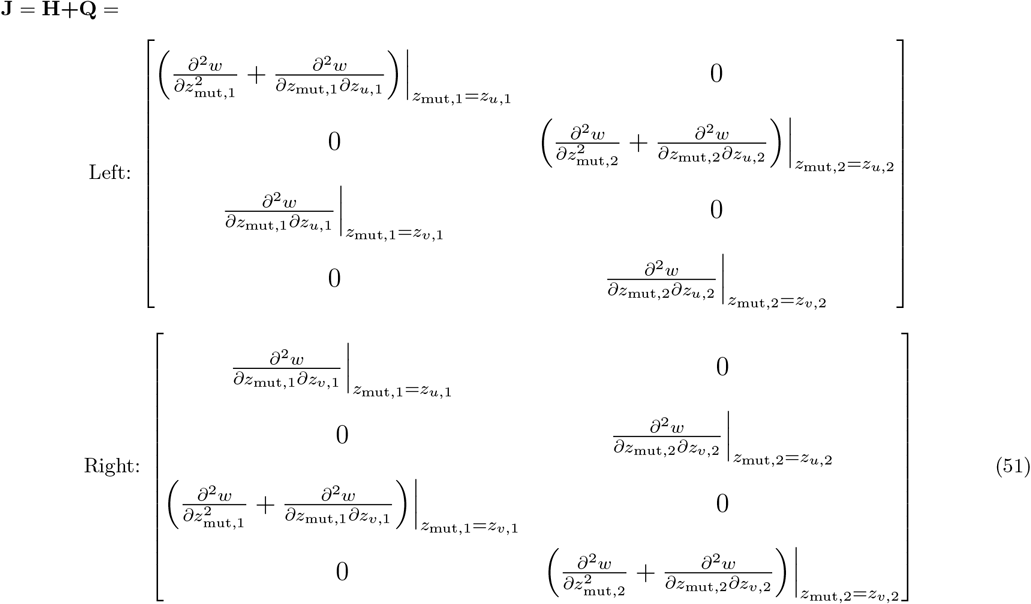

## D Analysis with payoffs of the form *π* = *B · C*

### D.1 Fitness

The fitness *w* of a mutant with trait *z*_mut_ in a population with resident trait *z*_res_ is

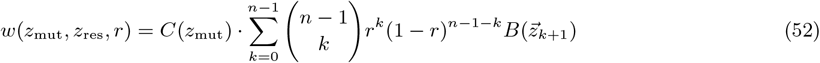

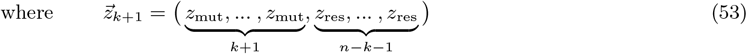

That is, 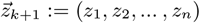. *z*_*i*_ = *z*_mut_ ∀*i* ≤ *k* + 1 and *z*_*i*_ = *z*_res_ ∀*i > k* + 1.

Note that

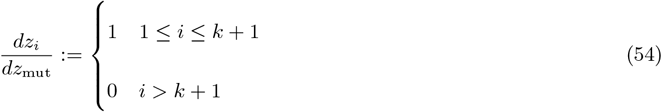

and 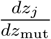 is defined analogously.

We assume that 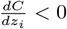 and 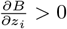 for any *i* ∈ {1, …, *n*}.

### D.2 Convergence stability

We derive the selection gradient as follows:

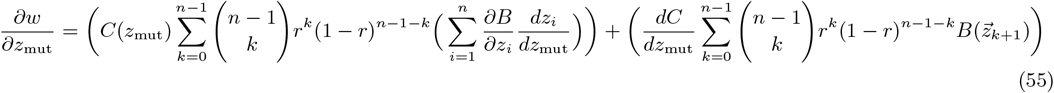

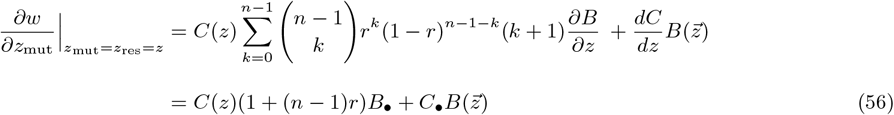

Where 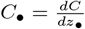 and 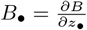 are the derivatives of *C* and *B* wrt. a focal investment (•) among the *n*, when all are evaluated at the resident point *z*. To derive equation (4), we assume exchangeability of investments evaluated at the same point.

The CS condition (inequality 14 in WL (2014)) is

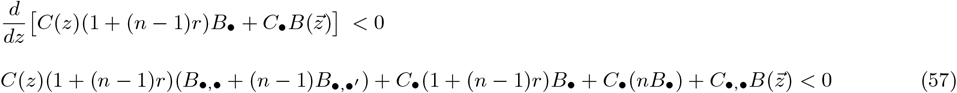

Where 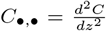 is the second derivative of the cost function, 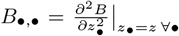 is the second derivative of the benefit function wrt. any focal trait, and 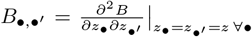 is the cross derivative of the benefit function wrt. focal trait *z*_•_ and any nonfocal trait 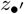, where all are evaluated at resident trait *z*.

### D.3 Evolutionary instability

Return to equation (3) above and take the second partial derivative of fitness wrt. *z*_mut_:

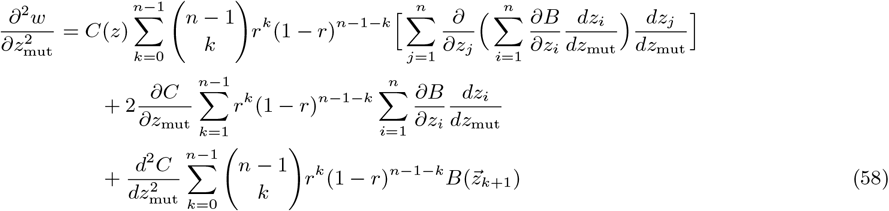

Now evaluate at a resident trait, such that *z*_mut_ = *z*_res_ = *z*:

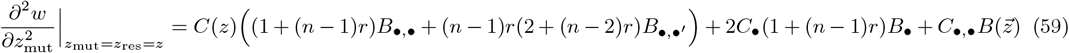

### D.4 Branching requires that the cross derivative is negative (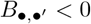)

From (6), we have

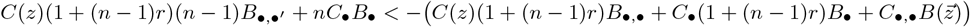

From (8), we have

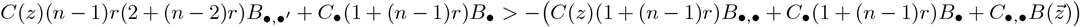

Taking the leftmost and rightmost terms, we have

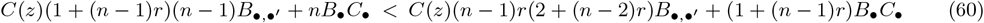

This implies 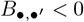 for branching, by the same logic as with *π* = *B − C*.

## E Proving properties of interchangeability

For clarity, write *B* = *B*(*z*_*i*_ (*x, y*), *z*_*j*_ (*x, y*)). From definition 1, we have:

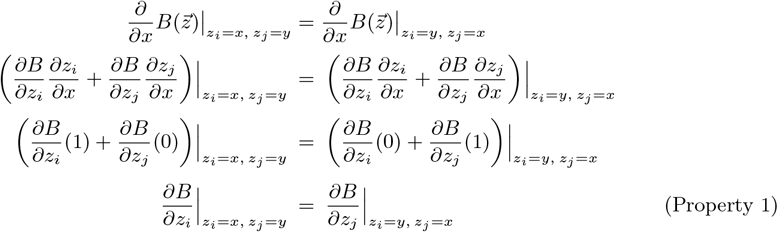

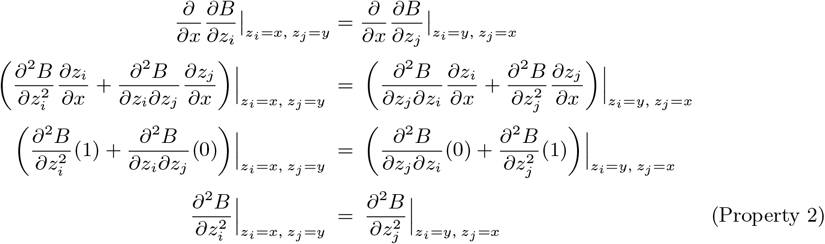

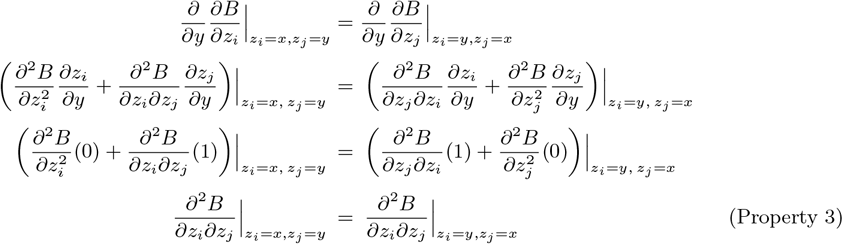

Properties 1-2 hold if the indices *i* and *j* are switched. This can be shown by applying the same derivation as above, taking 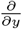 for property 1, then 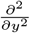 for property 2. Property 3 does not change if the indices are switched.

## F Proving uniqueness and dynamical stability of *p*_*\**_

### Result

*Let the payoff function for one public good for player i* ∈ (1, …, *n*) *be π*(*z*_*i*_, *·*) = *B*(*g*(*z*_*i*_, *·*)) *− C*(*z*_*i*_) *where the function* 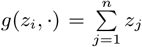 *(with other assumptions as defined in Methods), and let the investments of dimorphic types be (wlog) z*_*u*_ *> z*_*v*_ *≥* 0.

*If* 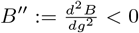 *on* ℝ^+^ *and interior solution p*_*\**_ ∈ (0, 1) *satisfying* 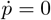 *exists, then p*_*\**_ *is unique and stable under replicator dynamics*.

**Proof:** We apply Result 3 of Peña et al. [2014a], which the authors extended in Peña et al. [2014b] to incorporate relatedness among individuals.

The focal individual plays with *n −* 1 others. Let *a*_*q*_ be the payoff to an individual investing *z*_*u*_ when *q* ≤ *n* – 1 others invest *z*_*u*_ (and *n −* 1 *− q* players invest *z*_*v*_), and let *b*_*q*_ be the payoff to an individual investing *z*_*v*_ when *q* others invest *z*_*u*_. Define *d*_*q*_ := *a*_*q*_ *− b*_*q*_ and denote Δ*d*_*q*_ = *d*_*q*+1_ *− d*_*q*_. Define *e*_*q*_ := *q*Δ*a*_*q−*1_ + (*n −* 1 *− q*)Δ*b*_*q*_, where Δ*a*_*q−*1_ = *a*_*q*_ *− a*_*q−*1_ and Δ*b*_*q*_ = *b*_*q*+1_ *− b*_*q*_. Finally, define *f*_*q*_ := *d*_*q*_ + *re*_*q*_. Denote Δ*f*_*q*_ = *f*_*q*+1_ *− f*_*q*_ and denote the gain sequence **f** := (*f*_0_, *f*_1_, …, *f*_*n−*1_).

These definitions follow Peña et al. [2014b] section 2.4, where we have used the subscript *q* instead of the authors’ *k* to distinguish from the subscript *k* (number of relatives) used in our definition of fitness. Under our simple relatedness scheme, the scaled relatedness coefficient *κ* from Peña et al. [2014b] is equal to the parameter *r*.

First, we expand *d*_*q*_ and *e*_*q*_ and derive *f*_*q*_ using the definition of payoff above:

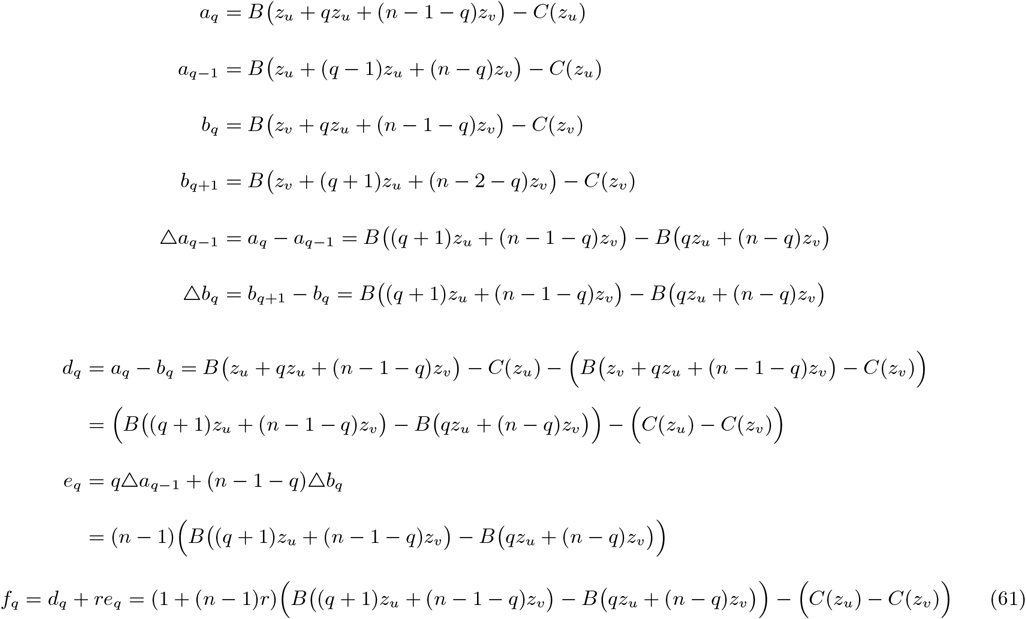

Following Peña et al. [2014b] equation (10), we can now write the gain function *G*:

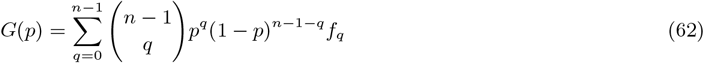

By Result 3 of Peña et al. [2014a], if the gain sequence **f** has a single sign change and *f*_0_ *>* 0, then any interior solution *p*_*\**_ ∈ (0, 1) satisfying 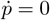 is unique and stable under replicator dynamics.

Note that we have assumed that public benefit *B* is a function of the sum of investments, denoting 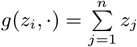 We now show that if 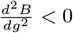 on ℝ^+^, then Δ*f*_*q*_ < 0.

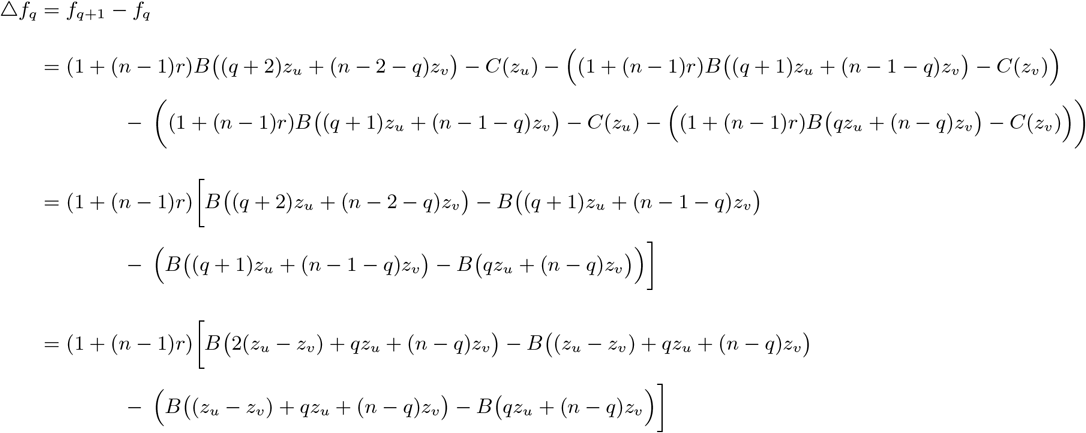

If *B*^*′′*^ *<* 0, the term in square brackets in the last line above is negative. This is because for any continuous, real-valued function *h*(*x*), for *x*_0_*< x*_1_ *< x*_2_ with 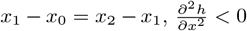 implies that *h*(*x*_2_) *− h*(*x*_1_) *< h*(*x*_1_) *− h*(*x*_0_).

Also, the term (1 + (*n −* 1)*r*) is strictly positive. We conclude that Δ*f*_*q*_ *<* 0.

Lastly, to show that the gain sequence **f** has a single sign change given Δ*f*_*q*_ *<* 0, we must show that *f*_0_ *>* 0 and *f*_*n−*1_ *<* 0. Rewrite both of these expressions as follows:

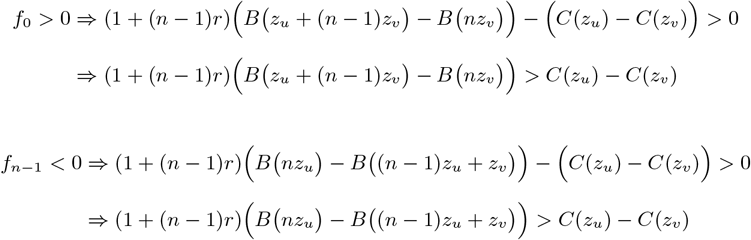

At a candidate solution *p*_*\**_ ∈ (0, 1) for which 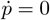, the following equality is true (see equations (40) and (41)):

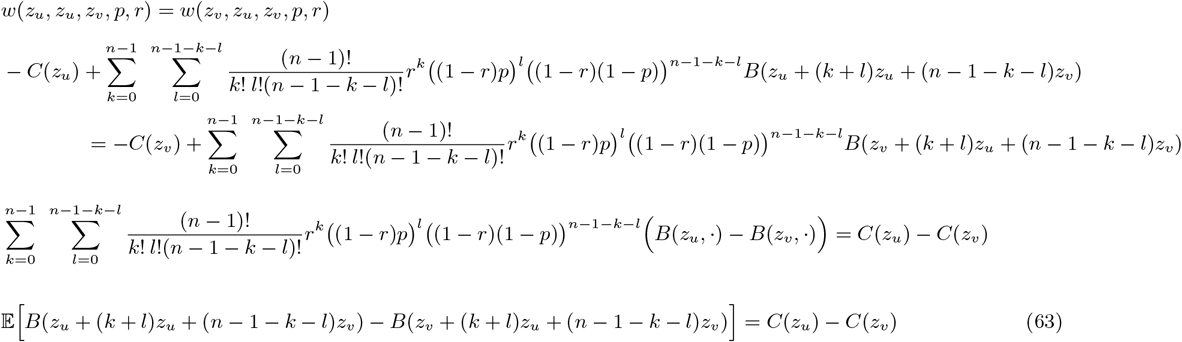

Show by contradiction that *B*^*′′*^ *<* 0 implies *f*_0_ *>* 0. Assume that *f*_0_ *<* 0. We have

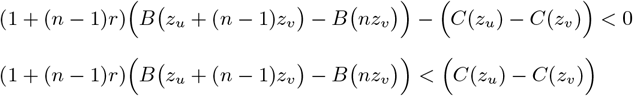

Because *B*^*′′*^ *<* 0, every value of *B* in the LHS expectation in equation (63) (i.e. with *q >* 0) is smaller than the interval *B z*_*u*_ + (*n −* 1)*z*_*v*_ *− B*(*nz*_*v*_). This implies

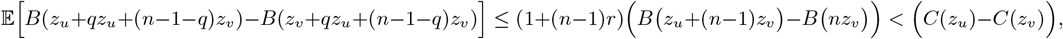

where *q* = *k* + *l*. This contradicts equation (63), so we conclude *f*_0_ *>* 0.

Similarly, assume that *f*_*n−*1_ *>* 0. We have

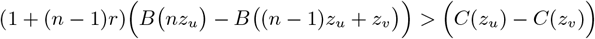

Because *B*^*′′*^ *<* 0, every value of *B* in the LHS expectation in equation (63) is larger than the interval *B(nz*_*u*_)*−B* ((*n −* 1)*z*_*u*_ + *z*_*v*_. This implies

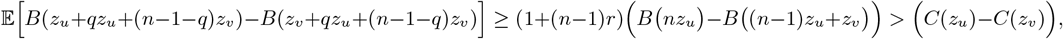

This again contradicts equation (63), so we conclude *f*_*n−*1_ *<* 0. ■

## Notes

### Competing Interest Statement

The authors have declared no competing interest.

