## Supplemental information for "The effects of group size and assortment on the evolution of division of labor"

### A Adaptive dynamics of investments in one public good

We derive the conditions for convergence stability and evolutionary instability of a monomorphic critical investment under a simple relatedness scheme. Our basic assumption is that fitness for a focal individual is given by the expected payoff from a public goods game with $n-1$ other players, each of whom is the same type as the focal individual with probability $r \in [0,1]$ (or is a randomly drawn type with probability $1-r$).

#### A.1 Interchangeable benefit functions

Define public benefit as a vector-valued function of $n$ investment traits, that is $B(\vec{z}) := B\big((z_1, z_2, \dots, z_n)\big)$.

**Definition 1.** For any $i, j \in (1, ..., n)$ s.t. $i \neq j$, let $z_i = x$ and $z_j = y$ where $x, y \in \mathbb{R}$. Investments are *interchangeable* iff. $B(\vec{z})\big|_{z_i=x,\, z_j=y} = B(\vec{z})\big|_{z_i=y,\, z_j=x}$

We can show that definition 1 implies the following equalities:

$$\frac{\partial B}{\partial z_i}\Big|_{z_i=x,\, z_j=y} = \frac{\partial B}{\partial z_j}\Big|_{z_i=y,\, z_j=x} \tag{13}$$

$$\frac{\partial^2 B}{\partial z_i^2}\Big|_{z_i=x,\, z_j=y} = \frac{\partial^2 B}{\partial z_j^2}\Big|_{z_i=y,\, z_j=x} \tag{14}$$

$$\frac{\partial^2 B}{\partial z_i \partial z_j}\Big|_{z_i=x,\, z_j=y} = \frac{\partial^2 B}{\partial z_i \partial z_j}\Big|_{z_i=y,\, z_j=x} \tag{15}$$

That is, if the investments of two individuals among the $n$ switch, the first and second partial derivatives of $B$ with respect to their investments also switch. We use the definition of interchangeability to define fitness (16). We use the first interchangeability property ((13)) above to derive the selection gradient (18). And, we use the latter properties ((14)) and ((15)) above to derive the convergence stability condition (19) and the evolutionary instability condition (22).

#### A.2 Fitness

The fitness $w$ of a mutant with trait $z_{\text{mut}}$ in a population with resident trait $z_{\text{res}}$ is

$$w(z_{\text{mut}}, z_{\text{res}}, r) = -C(z_{\text{mut}}) + \sum_{k=0}^{n-1} \binom{n-1}{k} r^k (1-r)^{n-1-k} B(\vec{z}_{k+1}) \tag{16}$$

$$\text{where} \qquad \vec{z}_{k+1} = \big(\underbrace{z_{\text{mut}}, \dots, z_{\text{mut}}}_{k+1}, \underbrace{z_{\text{res}}, \dots, z_{\text{res}}}_{n-k-1}\big)$$

That is, $\vec{z}_{k+1} := (z_1, z_2, \dots, z_n)$ st. $z_i = z_{\text{mut}}\ \forall i \leq k+1$ and $z_i = z_{\text{res}}\ \forall i > k+1$.

Note that

$$\frac{dz_i}{dz_{\text{mut}}} := \begin{cases} 1 & 1 \leq i \leq k+1 \\ 0 & i > k+1 \end{cases}$$

and $\frac{dz_j}{dz_{\text{mut}}}$ is defined analogously.

### B Evolutionary branching conditions

#### B.1 Convergence stability

We derive the selection gradient as follows:

$$\frac{\partial w}{\partial z_{\mathrm{mut}}} = -\frac{dC}{dz_{\mathrm{mut}}} + \sum_{k=0}^{n-1} \binom{n-1}{k} r^k (1-r)^{n-1-k} \Big( \sum_{i=1}^{n} \frac{\partial B}{\partial z_i} \frac{dz_i}{dz_{\mathrm{mut}}} \Big) \tag{17}$$

$$\begin{aligned} \frac{\partial w}{\partial z_{\mathrm{mut}}}\Big|_{z_{\mathrm{mut}}=z_{\mathrm{res}}=z} &= -\frac{dC}{dz} + \sum_{k=0}^{n-1} \binom{n-1}{k} r^k (1-r)^{n-1-k} (k+1) \frac{\partial B}{\partial z} \\ &= -C_\bullet + (1+(n-1)r) B_\bullet \end{aligned} \tag{18}$$

Where $C_\bullet = \frac{dC}{dz_\bullet}$ and $B_\bullet = \frac{\partial B}{\partial z_\bullet}$ are the derivatives of $C$ and $B$ wrt. a focal investment ($\bullet$) among the $n$, when all are evaluated at the resident point $z$. Note that the derivation of mutant fitness (18) assumes interchageability of investments as defined above.

The CS condition is

$$\begin{aligned} &\frac{d}{dz}\big[ -C_\bullet + (1+(n-1)r)B_\bullet \big] \; < 0 \\ &-\frac{d^2C}{dz^2} + (1+(n-1)r)\frac{d}{dz}\Big[\frac{\partial B}{\partial z_\bullet}\Big|_{z_\bullet = z\ \forall \bullet}\Big] \; < 0 \\ &-C_{\bullet,\bullet} + (1+(n-1)r)\big(B_{\bullet,\bullet} + (n-1)B_{\bullet,\bullet'}\big) \; < 0 \end{aligned} \tag{19}$$

Where $C_{\bullet,\bullet} = \frac{d^2C}{dz^2}$ is the second derivative of the cost function, $B_{\bullet,\bullet} = \frac{\partial^2 B}{\partial z_\bullet^2}\big|_{z_\bullet = z\ \forall \bullet}$ is the second derivative of the benefit function wrt. any focal trait, and $B_{\bullet,\bullet'} = \frac{\partial^2 B}{\partial z_\bullet \partial z_{\bullet'}}\big|_{z_\bullet = z_{\bullet'} = z\ \forall \bullet}$ is the cross-derivative of the benefit function wrt. focal trait $z_\bullet$ and any nonfocal trait $z_{\bullet'}$, where all are evaluated at resident trait $z$. Note that (19) is equivalent to inequality (14) in Wakano and Lehmann [2014].

#### B.2 Evolutionary instability

Return to equation (17) above and take the second partial derivative of fitness wrt. $z_{\mathrm{mut}}$:

$$\begin{aligned} \frac{\partial^2 w}{\partial z_{\mathrm{mut}}^2} &= -\frac{\partial^2 C}{\partial z_{\mathrm{mut}}^2} + \sum_{k=0}^{n-1} \binom{n-1}{k} r^k (1-r)^{n-1-k} \Big[ \sum_{j=1}^{n} \frac{\partial}{\partial z_j} \Big( \sum_{i=1}^{n} \frac{\partial B}{\partial z_i} \frac{dz_i}{dz_{\mathrm{mut}}} \Big) \frac{dz_j}{dz_{\mathrm{mut}}} \Big] \\ &= -\frac{\partial^2 C}{\partial z_{\mathrm{mut}}^2} + \sum_{k=0}^{n-1} \binom{n-1}{k} r^k (1-r)^{n-1-k} \Big[ \sum_{j=1}^{n} \sum_{i=1}^{n} \frac{\partial^2 B}{\partial z_i \partial z_j} \frac{dz_i}{dz_{\mathrm{mut}}} \frac{dz_j}{dz_{\mathrm{mut}}} \Big] \end{aligned} \tag{20}$$

Now evaluate at a resident trait, such that $z_{\text{mut}} = z_{\text{res}} = z$:

$$\begin{aligned}
\frac{\partial^2 w}{\partial z_{\text{mut}}^2}\Big|_{z_{\text{mut}}=z_{\text{res}}=z} &= \left( -\frac{\partial^2 C}{\partial z_{\text{mut}}^2} + \sum_{k=0}^{n-1} \binom{n-1}{k} r^k (1-r)^{n-1-k} \Big[ \sum_{j=1}^{n} \sum_{i=1}^{n} \frac{\partial^2 B}{\partial z_i \partial z_j} \frac{dz_i}{dz_{\text{mut}}} \frac{dz_j}{dz_{\text{mut}}} \Big] \right)\Big|_{z_{\text{mut}}=z_{\text{res}}=z} \\
&= -\frac{d^2 C}{dz^2} + \sum_{k=0}^{n-1} \binom{n-1}{k} r^k (1-r)^{n-1-k} \left( (k+1)\frac{\partial^2 B}{\partial z_\bullet^2} + (k+1)k \frac{\partial^2 B}{\partial z_\bullet \partial z_{\bullet'}} \right) \\
&= -C_{\bullet,\bullet} + (1+(n-1)r)B_{\bullet,\bullet} + \big((n-1)r(2+(n-2)r)\big)B_{\bullet,\bullet'} \qquad (21)
\end{aligned}$$

Where $z_\bullet$ and $z_{\bullet'}$ are generic focal and nonfocal traits, as defined above, and we evaluated the sum in Mathematica. After simplifying, we obtain the following evolutionary instability condition:

$$-C_{\bullet,\bullet} + B_{\bullet,\bullet} + (n-1)r\big(B_{\bullet,\bullet} + 2B_{\bullet,\bullet'}\big) + (n-1)(n-2)r^2 B_{\bullet,\bullet'} > 0 \qquad (22)$$

This is equivalent to inequality (29) in Wakano and Lehmann [2014], with $\triangle w = \frac{\partial^2 w}{\partial z_{\text{mut}}^2}\Big|_{z_{\text{mut}}=z_{\text{res}}=z}$, $w_{SS} = B_{\bullet,\bullet} - C_{\bullet,\bullet}$, $w_{SD} = 2(n-1)B_{\bullet,\bullet'}$, $w_{DD} = (n-1)B_{\bullet,\bullet}$, $w_{DD'} = (n-1)(n-2)B_{\bullet,\bullet'}$, $R_2 = r$, $R_3 = r^2$, and $\triangle r = 0$ by assumption.

##### B.3 Branching requires that the cross derivative is negative ($B_{\bullet,\bullet'} < 0$) and relatedness is nonclonal ($r < 1$)

From CS condition (19) above, we have

$$(n-1)(1+(n-1)r)B_{\bullet,\bullet'} < C_{\bullet,\bullet} - (1+(n-1)r)B_{\bullet,\bullet}$$

And from E(in)S condition (22) above, we have

$$\begin{aligned}
2(n-1)rB_{\bullet,\bullet'} + (n-1)(n-2)r^2 B_{\bullet,\bullet'} &> C_{\bullet,\bullet} - (1+(n-1)r)B_{\bullet,\bullet} \\
\Rightarrow\ (n-1)r(2+(n-2)r)B_{\bullet,\bullet'} &> C_{\bullet,\bullet} - (1+(n-1)r)B_{\bullet,\bullet}
\end{aligned}$$

Combining these, we obtain

$$(n-1)(1+(n-1)r)B_{\bullet,\bullet'} \ < \ C_{\bullet,\bullet} - (1+(n-1)r)B_{\bullet,\bullet} \ < \ (n-1)r(2+(n-2)r)B_{\bullet,\bullet'} \qquad (23)$$

When $r = 0$, we obtain the standard evolutionary branching conditions with no assortment, which imply that costs, focal benefits, and cross-benefits must all be saturating, that is $C_{\bullet,\bullet} < 0$, $B_{\bullet,\bullet} < 0$, $B_{\bullet,\bullet'} < 0$.

When $r \in (0,1)$, we can easily show that $(n-1)(1+(n-1)r) > (n-1)r(2+(n-2)r)$, implying that the cross-derivative must again be negative ($B_{\bullet,\bullet'} < 0$) for evolutionary branching conditions to be met (from the left and right sides of (23)). When $r = 1$, we obtain $n(n-1)B_{\bullet,\bullet'} < C_{\bullet,\bullet} - B_{\bullet,\bullet} < n(n-1)B_{\bullet,\bullet'}$ which is false, so evolutionary branching cannot happen with $r = 1$.

#### B.4 General class of public benefit functions for which branching implies diminishing $B$, $C$

Our basic inference from the general branching conditions (23) is that $B_{\bullet,\bullet'} < 0$. For additive or combinatorial public benefit functions (defined below), we can further show that $B_{\bullet,\bullet} \leq B_{\bullet,\bullet'}$, which in turn implies that $B_{\bullet,\bullet}, C_{\bullet,\bullet} < 0$. In this section, we define the generic properties of $B$ that allow us to make this inference.

Let $B : f(\vec{z}) \to \mathbb{R}^+$, where we assume $\frac{dB}{df} > 0$ and $\forall j \in (1, ..., n)$, $\frac{\partial f}{\partial z_j} > 0$, i.e. the effect on $B$ of each individual's investment is positive. We have

$$\frac{\partial B}{\partial z_i} = \frac{\partial f}{\partial z_i}\frac{dB}{dz_i}$$
$$\frac{\partial^2 B}{\partial z_i^2} = \Big(\frac{\partial f}{\partial z_i}\Big)^2\frac{d^2 B}{df^2} + \frac{\partial^2 f}{\partial z_i^2}\frac{dB}{df}$$
$$\frac{\partial^2 B}{\partial z_i^2} = \frac{\partial f}{\partial z_i}\frac{\partial f}{\partial z_j}\frac{d^2 B}{df^2} + \frac{\partial^2 f}{\partial z_i \partial z_j}\frac{dB}{df}$$

The latter two terms, evaluated at the monomorphic critical point, are $B_{\bullet,\bullet}$ and $B_{\bullet,\bullet'}$.

Note that $\left(\frac{\partial f}{\partial z_i}\right)^2 = \frac{\partial f}{\partial z_i}\frac{\partial f}{\partial z_j}$ due to interchangeability of investments, and $\frac{dB}{df} > 0$ by assumption.

Therefore, if $\frac{\partial^2 f}{\partial z_i^2} \leq \frac{\partial^2 f}{\partial z_i \partial z_j}$, then $B_{\bullet,\bullet} \leq B_{\bullet,\bullet'}$. The standard additive function $f : \sum_{j=1}^{n} z_j \to \mathbb{R}$ satisfies this requirement. More generally, any function $f$ that is a sum of products of all combinations of $k$ investments, or a sum of these sums for multiple $k$ values, satisfies this requirement (where $k \in \{1, ..., n\}$). If $k = 1$, then $f$ is the canonical function above; if $k = n$, then $f$ is the product of all $n$ investments; and for example, if $k = 2$, then $f$ is the sum of all pairs of investments, that is $f : \sum_{i \neq j} z_i z_j \to \mathbb{R}$.

We can construct a counterexample $f$ function for which investments are interchangeable and yet $\frac{\partial^2 f}{\partial z_i^2} > \frac{\partial^2 f}{\partial z_i \partial z_j}$: Let $f(\vec{z}) = \sum_{j=1}^{n} z_j^{\alpha}$. Then for any $i$ and $j$ from 1 to $n$ with $i \neq j$, we have $\frac{\partial^2 f}{\partial z_i^2} = \alpha(\alpha - 1)z_i^{\alpha-2}$, and $\frac{\partial^2 f}{\partial z_i \partial z_j} = 0$. Therefore if $\alpha > 1$, then $\frac{\partial^2 f}{\partial z_i^2} > \frac{\partial^2 f}{\partial z_i \partial z_j}$, and evolutionary branching does not imply $B_{\bullet,\bullet'}, B_{\bullet,\bullet}, C_{\bullet,\bullet} < 0$. If $0 \leq \alpha \leq 1$, then $\frac{\partial^2 f}{\partial z_i^2} \leq \frac{\partial^2 f}{\partial z_i \partial z_j}$, so evolutionary branching implies $B_{\bullet,\bullet'}, B_{\bullet,\bullet}, C_{\bullet,\bullet} < 0$.

#### B.5 Other observations

By contrast, if the monomorphic critical point is convergently unstable and evolutionarily unstable, we cannot infer the sign of $B_{\bullet,\bullet'}$ or the signs of $B_{\bullet,\bullet}$ of $C_{\bullet,\bullet}$. This is because in the unstable setting, the conditions for the second partial derivatives can be found by moving *both* the LHS and RHS of branching conditions (23) to the right (left) of the middle term. Under these different conditions, there is no information about the order of the LHS and RHS, which under evolutionary branching conditions (23) implies that $B_{\bullet,\bullet'} < 0$ and therefore $B_{\bullet,\bullet}$, $C_{\bullet,\bullet} < 0$.

The same analysis applies for multiplicative payoff functions of the form $\pi_i = B \cdot C$ (see S.I. D below). We can show that if public benefits are in the additive/combinatorial family of functions, then $B_{\bullet,\bullet'}$, $B_{\bullet,\bullet} < 0$ are necessary for evolutionary branching to occur.

##### B.6 Effect of assortment $r$ on critical point $z_*$

The conditions for evolutionary branching imply that $\frac{\partial z_*}{\partial r} > 0$, that is, higher levels of assortment increase the critical level.

Proof: The monomorphic critical point $z_*$ can be found by solving

$$(1 + (n-1)r)B_\bullet - C_\bullet = 0 \tag{24}$$

for resident trait level $z$. Write $z_* := z_*(n, r)$ as a function of $n$ and $r$, and at that point, write $B_\bullet := B_\bullet(z_*, ..., z_*)$ and $C_\bullet := C_\bullet(z_*)$ as functions of $z_*$. Now take the total derivative of (24) with respect to $r$:

$$\begin{aligned}
&\frac{\partial}{\partial r}\big[(1 + (n-1)r)B_\bullet - C_\bullet\big] = 0 \\
&(n-1)B_\bullet + (1 + (n-1)r)\big(B_{\bullet,\bullet}\frac{\partial z_*}{\partial r} + (n-1)B_{\bullet,\bullet'}\frac{\partial z_*}{\partial r}\big) - C_{\bullet,\bullet}\frac{\partial z_*}{\partial r} = 0 \\
&\frac{\partial z_*}{\partial r}\Big((1 + (n-1)r)(B_{\bullet,\bullet} + (n-1)B_{\bullet,\bullet'}) - C_{\bullet,\bullet}\Big) = -(n-1)B_\bullet \\
&\frac{\partial z_*}{\partial r} = \frac{-(n-1)B_\bullet}{(1 + (n-1)r)(B_{\bullet,\bullet} + (n-1)B_{\bullet,\bullet'}) - C_{\bullet,\bullet})}
\end{aligned} \tag{25}$$

The numerator in (25) is negative because we assume $B_\bullet > 0$, and the denominator is negative under the general branching conditions (23). This implies that $\boxed{\frac{\partial z_*}{\partial r} > 0}$.

##### B.7 Branching conditions when $B$ is a function of additive investments

If $B$ is a function of the sum of investments $\sum z_j$, then $B_{\bullet,\bullet} = B_{\bullet,\bullet'}$. In this case, we have

$$n(1 + (n-1)r)B_{\bullet,\bullet} < C_{\bullet,\bullet} < \Big((1 + (n-1)r) + (n-1)r(2 + (n-2)r)\Big)B_{\bullet,\bullet} \tag{26}$$

Because $1 + (n-1)r \;>\; 2r + (n-2)r^2$, the RHS of (26) is greater than the LHS *iff.* $B_{\bullet,\bullet} < 0$, which further implies that $C_{\bullet,\bullet} < 0$ in this case. We can divide by $B_{\bullet,\bullet}$ to obtain

$$n(1 + (n-1)r) > \frac{C_{\bullet,\bullet}}{B_{\bullet,\bullet}} > (1 + (n-1)r) + (n-1)r(2 + (n-2)r) \tag{27}$$

This pair of inequalities is used in the following section.

#### B.8 Analysis with numerical payoff functions

##### B.8.1 Model 1

Let $\pi_i = (\sum_{j=1}^{n} z_j)^{a_1} - z_i^{a_2}$. We have

$$\frac{\partial B}{\partial z_i} = a_1(\sum_{j=1}^{n} z_j)^{a_1-1} \Rightarrow B_\bullet = a_1(nz)^{a_1-1}$$
$$\frac{\partial^2 B}{\partial z_i^2} = a_1(a_1-1)(\sum_{j=1}^{n} z_j)^{a_1-2} \Rightarrow B_{\bullet,\bullet} = a_1(a_1-1)(nz)^{a_1-2}$$
$$\frac{\partial C}{\partial z_i} = C_\bullet = a_2 z^{a_2-1}$$
$$\frac{\partial^2 C}{\partial z_i^2} = C_{\bullet,\bullet} = a_2(a_2-1)z^{a_2-2}$$

We solve for the critical point $z_*$ by setting $(1+(n-1)r)B_\bullet - C_\bullet = 0$ and solving for $z$, to obtain

$$z_* = \Big((1+(n-1)r)n^{a_1-1}\frac{a_1}{a_2}\Big)^{\frac{1}{a_2-a_1}} \tag{28}$$

Note that

$$\frac{\partial z_*}{\partial r} = (n-1)\Big((1+(n-1)r)n^{a_1-1}\frac{a_1}{a_2}\Big)^{\frac{1}{a_2-a_1}-1} \tag{29}$$

Equation (29) and our baseline assumptions that $a_1, a_2 > 0$ imply that $\frac{\partial z_*}{\partial r} > 0$.

Plug the expressions for $B_{\bullet,\bullet}$ and $C_{\bullet,\bullet}$ into (27) to obtain evolutionary branching conditions for model 1:

$$n(1+(n-1)r) > \frac{a_2(a_2-1)z_*^{a_2-2}}{a_1(a_1-1)(nz_*)^{a_1-2}} > (1+(n-1)r) + (n-1)r(2+(n-2)r)$$
$$n(1+(n-1)r) > \frac{a_2(a_2-1)}{a_1(a_1-1)n^{a_1-2}}z_*^{a_2-a_1} > (1+(n-1)r) + (n-1)r(2+(n-2)r)$$
$$n(1+(n-1)r) > n(1+(n-1)r)\,\frac{a_2-1}{a_1-1} > (1+(n-1)r) + (n-1)r(2+(n-2)r)$$
$$1 > \frac{a_2-1}{a_1-1} > \frac{1}{n} + \Big(\frac{n-1}{n}\Big)\frac{r(2+(n-2)r)}{(1+(n-1)r)} \tag{30}$$

As $r$ increases, the region of $(a_1, a_2)$-space that allows evolutionary branching becomes smaller. The area of this region is directly proportional to the quantity

$$V \;=\; \frac{n-1}{n}\Big(1 - \frac{r(2+(n-2)r)}{(1+(n-1)r)}\Big) \tag{31}$$

We can show that $V$ increases with $n$ by evaluating the quantity

$$\triangle_n V = V_{n+1} - V_n = \frac{(1-r)(1+(n-1)2r(1+nr))}{n(n+1)(1+(n-1)r)(1+nr)} \tag{32}$$

Each term in this quotient is positive, so $\triangle_n V > 0$.

The rate of change of $V$ with respect to $r$ is

$$\frac{\partial V}{\partial r} = -\frac{n-1}{n}\left(\frac{2+2(n-2)r+(n-1)(n-2)r^2}{1+2(n-1)r+(n-1)^2r^2}\right) \tag{33}$$

The term in parentheses is strictly positive, so $\frac{\partial V}{\partial r} < 0$.

Finally, we can examine the cross-effects of $n$ and $r$ by computing the following

$$\frac{\partial}{\partial r}\triangle_n V \;=\; \frac{\partial V_{n+1}}{\partial r} - \frac{\partial V_n}{\partial r} \;=\; \frac{1}{(1+(n-1)r)^2} - \frac{1}{(1+nr)^2} - \frac{2}{n(n+1)} \tag{34}$$

This expression is typically negative, meaning that for a given $r$, if $n$ increases then the effect on $V$ of increasing $r$ will become more negative. However, for $n \geq 6$, there is a range of small $r$ values for which the expression is positive, meaning that increasing $n$ dampens the negative effect of $r$ on $V$.

###### B.8.2 Model 2

Let $\pi_i = \big[-a_1\big(\sum_{j=1}^{n} z_j\big)^2 + a_2\big(\sum_{j=1}^{n} z_j\big)\big] - \big[-a_3 z_i^2 + a_4 z_i\big]$. The first term in square brackets is public benefit, and the second (subtracted) bracketed term is private cost. We have

$$\begin{aligned}
&\frac{\partial B}{\partial z_i} = -2a_1\sum_{j=1}^{n} z_j + a_2 \;\Rightarrow\; B_\bullet = -2a_1 nz + a_2\\
&\frac{\partial^2 B}{\partial z_i^2} = B_{\bullet,\bullet} = -2a_1\\
&\frac{dC}{dz_i} = -2a_3 z_i + a_4 \;\Rightarrow\; C_\bullet = -2a_3 z + a_4\\
&\frac{d^2 C}{dz_i^2} = C_{\bullet,\bullet} = -2a_3
\end{aligned}$$

From equation (27), the evolutionary branching conditions for model 2 are as follows:

$$n(1+(n-1)r) > \frac{a_3}{a_1} > 1+(n-1)r+(n-1)r(2+(n-2)r) \tag{35}$$

The region of $(a_1, a_3)$-space that allows evolutionary branching is a triangle in the upper-right quadrant that is defined by the LHS and RHS of (35). The area of the region is directly proportional to the difference $D$ between these two quantities:

$$\begin{aligned}
D &= n(1+(n-1)r) - (1+(n-1)r) - (n-1)r(2+(n-2)r)\\
&= (n-1)\big(1+(n-1)r - r(2+(n-2)r)\big)\\
&= (n-1)\big(1+(n-3)r-(n-2)r^2\big)
\end{aligned} \tag{36}$$

Show that $D$ increases with $n$ (i.e. $\triangle_n D > 0$):

$$\triangle_n D = D_{n+1} - D_n = (1-r)(1+2(n-1)r) \tag{37}$$

And the rate of change of $V$ wrt. $r$ is

$$\begin{aligned}\frac{\partial D}{\partial r} &= (n-1)(n-3-2(n-2)r) \\ &= (n-1)(n(1-2r)-(3-4r))\end{aligned} \tag{38}$$

With $n=2$, $\frac{\partial V}{\partial r}=-1$; with $n=3$, $\frac{\partial V}{\partial r}=0$ for $r=0$ and decreases linearly with $r$; and with $n\geq 4$, $\frac{\partial V}{\partial r}=0$ at $r_*=\left(\frac{1}{2}\right)\left(\frac{n-3}{n-2}\right)$ and decreases linearly with $r$, meaning that $\frac{\partial V}{\partial r}>0$ for $r<r_*$ and $\frac{\partial V}{\partial r}<0$ for $r>r_*$. The cross-effects of $n$ and $r$ are given by

$$\frac{\partial}{\partial r}\triangle_n D = \frac{\partial D_{n+1}}{\partial r} - \frac{\partial D_n}{\partial r} \; = \; 2n(1-2r)-(3-4r) \tag{39}$$

Which are linearly decreasing with $r$, similar to $\frac{\partial D}{\partial r}$.

##### B.9 Adaptive dynamics with two types, with one public good and assortment ($r>0$)

After evolutionary branching, there are two resident types, $u$ and $v$, with traits $z_u$ ($z_v$) and frequencies $p\,(1-p)$, where $p\in(0,1)$.

Consider a mutant with investment $z_{\text{mut}}$ in a resident population with investments $(z_u,z_v)$. We can write the payoff to the mutant from the public goods game as

$$\pi = B(\vec{z}_{k+1,l}) - C(z_{\text{mut}})$$

where $B(\vec{z}_{k+1,l})$ is a vector of which $k+1$ elements are equal to $z_{\text{mut}}$, while $l$ elements are equal to $z_u$ and $n-1-k-l$ elements are equal to $z_v$.

The fitness $w$ of a mutant with trait $z_{\text{mut}}$ in a population with resident traits $(z_u,z_v)$ is

$$\begin{aligned} w(z_{\text{mut}}, z_u, z_v, p, r) &= -C(z_{\text{mut}}) + \sum_{k=0}^{n-1}\binom{n-1}{k} r^k (1-r)^{n-1-k} \sum_{l=0}^{n-1-k}\binom{n-1-k}{l} p^l (1-p)^{n-1-k-l}\, B(\vec{z}_{k+1,l}) \\ &= -C(z_{\text{mut}}) + \sum_{k=0}^{n-1}\sum_{l=0}^{n-1-k-l}\binom{n-1}{k}\binom{n-1-k}{l} r^k (1-r)^{n-1-k-l+l} p^l (1-p)^{n-1-k-l} B(\vec{z}_{k+1,l}) \\ &= -C(z_{\text{mut}}) + \sum_{k=0}^{n-1}\sum_{l=0}^{n-1-k-l}\frac{(n-1)!}{k!\,l!(n-1-k-l)!} r^k \big((1-r)p\big)^l \big((1-r)(1-p)\big)^{n-1-k-l} B(\vec{z}_{k+1,l}) \end{aligned} \tag{40}$$

Thus, the public benefit $B(\vec{z}_{k+1,l})$ that the mutant receives from the game has a trinomial distribution. For each nonfocal player $j$, with probability $r$, $j$ has trait $z_{\text{mut}}$; with probability $(1-r)p$, has trait $z_u$; and with probability $(1-r)(1-p)$, has trait $z_v$.

The fitness of a type $u$ individual in the resident population is written $w(z_u,z_u,z_v,p,r)$, i.e. treating the focal 'mutant' as a type $u$ individual in (40) (fitness of type $v$ is defined analogously).

The dynamics of $p$ (the frequency of type $u$) are given by

$$\dot{p} = p(1-p)\big(w(z_u, z_u, z_v, p, r) - w(z_v, z_u, z_v, p, r)\big) \tag{41}$$

For the summed-investment family of payoff functions, if an internal equilibrium frequency $p_* \in (0,1)$ satisfying $\dot{p} = 0$ exists, then it is dynamically stable and unique. We prove this below in S.I. F. This allows us to initialize dimorphic investments $z_u$ and $z_v$ close to a branching point, where they have a unique frequency equilibrium $p_*(z_u, z_v)$, and derive their adaptive dynamics by integrating the two-type selection gradient along the manifold defined by $p_*$.

Note: if we distinguished relatedness within resident types ($r_{\text{type}}$) from that within focal strains ($r_{\text{mut}}$), we could compute the resulting adaptive dynamics by substituting $r_{\text{type}}$ into (41) in our solution for $p_*$, and substituting $r_{\text{mut}}$ in our equation for mutant fitness (40).

The two-type selection gradient is computed by taking the partial derivative of fitness with respect to the mutant trait and evaluating at $z_{\text{mut}} = z_u$ (for type $u$) and $z_{\text{mut}} = z_v$ (for type $v$) when the resident traits $z_u$ ($z_v$) are at equilibrium frequencies $p_*$ $(1-p_*)$:

$$(\dot{z_u}, \dot{z_v}) \propto \left( \frac{\partial w}{\partial z_{\text{mut}}}\Big|_{z_{\text{mut}}=z_u}, \frac{\partial w}{\partial z_{\text{mut}}}\Big|_{z_{\text{mut}}=z_v} \right)\Big|_{\vec{z}_{\text{res}}=(z_u, z_v)\,;\, p=p_*} \tag{42}$$

We derive an adaptive dynamical trajectory of $z_u$ and $z_v$ by choosing initial traits $(z_{u0}, z_{v0})$ with internal frequency equilibrium $p_{*0}$ and integrating the two-type selection gradient (42). In every such case, if $z_{u0} > z_{v0}$, then $z_u$ evolves upward and $z_v$ downward, and vice versa. The resident traits may evolve to a dimorphic critical point, at which $(\dot{z}_u, \dot{z}_v) = (0,0)$, or to a point at which $p_* = 0$ or 1, so one type fixes and the population reverts to a monomorphic state.

Finally, to analyze the convergence and evolutionary (in)stability of a dimorphic critical point, we define the Hessian and Jacobian matrices of the selection gradient, following Leimar [2009]:

$$\mathbf{H} = \begin{bmatrix} \frac{\partial^2 w}{\partial z_{\text{mut}}^2}\Big|_{z_{\text{mut}}=z_u} & 0 \\ 0 & \frac{\partial^2 w}{\partial z_{\text{mut}}^2}\Big|_{z_{\text{mut}}=z_v} \end{bmatrix} \tag{43}$$

$$\mathbf{J} = \begin{bmatrix} \left(\frac{\partial^2 w}{\partial z_{\text{mut}}^2} + \frac{\partial^2 w}{\partial z_{\text{mut}} \partial z_u}\right)\Big|_{z_{\text{mut}}=z_u} & 0 \\ 0 & \left(\frac{\partial^2 w}{\partial z_{\text{mut}}^2} + \frac{\partial^2 w}{\partial z_{\text{mut}} \partial z_v}\right)\Big|_{z_{\text{mut}}=z_v} \end{bmatrix} \tag{44}$$

Where all elements of both matrices are evaluated at the resident dimorphic point $(z_u, z_v, p_*)$.

If the matrix J is negative definite, then the dimorphism is convergence stable. If the matrix H is negative definite, it is evolutionarily stable. If H is positive definite or indefinite, further evolutionary branching into additional types will occur. It is helpful that the definiteness of either H or J depends entirely on the signs of the diagonal elements, which are their eigenvalues.

For a given specification of payoff $\pi$, such as the power-function or quadratic forms analyzed above, we can check if the monomorphic critical point satisfies branching conditions, solve for a dimorphic critical point $(z_u, z_v)_*$ (if it exists) by setting (42) equal to 0, and determine the evolutionary and convergence (in)stability of $(z_u, z_v)_*$ by computing (43) and (44), respectively.

We depict the evolutionary stability properties of a large sample of dimorphic critical points in figure 3 (main text) for the power-function form of payoff. For very small values of $a_2 - a_1$, such that the rate at which cost diminishes only slightly exceeds the rate at which benefit diminishes, the critical investment for the lower-investing type decreases to an extremely small value on the order of $10^{-5}$. This causes the magnitudes of the components of the selection gradient for both types to increase rapidly. This in turn makes numerical integration of the dimorphic selection gradient challenging, because integrating that of the lower type for even a very small finite step size can cause the lower trait to drop below 0, which is undefined and breaks the solver that we used. That is the reason for the blank areas in the left part of the branching region in figure 3. Better algorithms could be used to solve for more critical points in this left region.

### C Adaptive dynamics with two public goods

With two public goods, each individual now has two investment traits, such that for individual $i \in (1, ..., n)$, their investments are written $\vec{z_i} = (z_{i,1}, z_{i,2})$.

The simplest way to define the payoff from two public goods is as follows. Let $B_k$ and $C_k$ denote the benefit and cost functions for good $k$, and let $\{z_{-i,k}\}$ denote the $n-1$ nonfocal investments in good $k$. Payoff for $i$ is

$$\pi\big(\vec{z_i}, \{z_{-i,1}\}, \{z_{-i,2}\}\big) = \underbrace{B_1\big(z_{i,1}, \{z_{-i,1}\}\big) - C_1(z_{i,1})}_{\text{Good 1}} + \underbrace{B_2\big(z_{i,2}, \{z_{-i,2}\}\big) - C_2(z_{i,2})}_{\text{Good 2}} \tag{45}$$

We label the two types that arise after branching $u$ and $v$, with investment traits $\vec{z}_u = (z_{u,1}, z_{u,2})$ and $\vec{z}_v = (z_{v,1}, z_{v,2})$ and frequencies $p$, $(1-p)$. As above, the fitness of a mutant with trait vector $\vec{z}_{\text{mut}}$ in this population is

$$w(\vec{z}_{\text{mut}}, \vec{z}_u, \vec{z}_v, p) = \mathbb{E}[\pi(z_{\text{mut}}, \cdot)|\vec{z_u}, \vec{z_v}, r, p] \tag{46}$$

The selection gradient $\big(\dot{z}_{u,1}, \dot{z}_{u,2}, \dot{z}_{v,1}, \dot{z}_{v,2}\big)$ for each of the four investment traits is proportional to

$$\left(\frac{\partial w}{\partial z_{\text{mut},1}}\Big|_{z_{\text{mut},1}=z_{u,1}}, \frac{\partial w}{\partial z_{\text{mut},2}}\Big|_{z_{\text{mut},2}=z_{u,2}}, \frac{\partial w}{\partial z_{\text{mut},1}}\Big|_{z_{\text{mut},1}=z_{v,1}}, \frac{\partial w}{\partial z_{\text{mut},2}}\Big|_{z_{\text{mut},2}=z_{v,2}}\right)\Big|_{\vec{z}_{\text{res}}=(\vec{z}_u, \vec{z}_v)\,;\,p=p_*} \tag{47}$$

Where $p = p_*$ is the solution of $w(\vec{z}_u, \vec{z}_u, \vec{z}_v, p, r) = w(\vec{z}_v, \vec{z}_u, \vec{z}_v, p, r)$ (see (41)).

If conditions (23) are true for either good 1 ($B_1$, $C_1$) or good 2 ($B_2$, $C_2$), then evolutionary branching into two types will occur. For payoff model 1, if (23) holds for both goods, the two types evolve to one of two dimorphic critical points. These points represent two characteristically different divisions of labor: exploitation (ex), in which some individuals invest more in both goods while others invest less in both; and specialization (sp), in which some individuals invest more in good 1 and less in good 2, while others invest more in good 2 and less in good 1. The exploitation and specialization outcomes are multistable. In our numerical investigations, the following basins for these outcomes hold for all parameters satisfying (23) (up to symmetry with $v$):

$$\text{At } t = 0,\ z_{u,1} > z_{v,1} \text{ and } z_{u,2} > z_{v,2} \Rightarrow \text{Exploitation } (u \text{ cooperates}) \tag{48}$$

$$\text{At } t = 0,\ z_{u,1} > z_{v,1} \text{ and } z_{u,2} < z_{v,2} \Rightarrow \text{Specialization } (u \text{ specializes in good 1}) \tag{49}$$

$$(\mathbf{H}_{kk})_{ij} = \frac{\partial^2 w}{\partial z_{\text{mut},i} \partial z_{\text{mut},j}} \Big|_{\vec{z}_{\text{mut}}=\vec{z}_k}$$

Where the first (second) partial derivative is taken with respect to a mutant of type $k$'s investment in public good $i$ ($j$), with $i, j \in \{1, 2\}$. Note that because in our payoff function (45) investments in different goods affect different summands (i.e. there are no interactions between goods), we know that $(\mathbf{H}_{kk})_{ij} = 0$ for $i \neq j$.

This gives us the following Hessian matrix for two goods and two types (using column ordering $(z_{u,1}, z_{u,2}, z_{v,1}, z_{v,2})$):

$$\mathbf{H} = \begin{bmatrix} \frac{\partial^2 w}{\partial z^2_{\text{mut},1}}\Big|_{z_{\text{mut},1}=z_{u,1}} & 0 & 0 & 0 \\ 0 & \frac{\partial^2 w}{\partial z^2_{\text{mut},2}}\Big|_{z_{\text{mut},2}=z_{u,2}} & 0 & 0 \\ 0 & 0 & \frac{\partial^2 w}{\partial z^2_{\text{mut},1}}\Big|_{z_{\text{mut},1}=z_{v,1}} & 0 \\ 0 & 0 & 0 & \frac{\partial^2 w}{\partial z^2_{\text{mut},2}}\Big|_{z_{\text{mut},2}=z_{v,2}} \end{bmatrix} \tag{50}$$

Where, as before with one good, all elements of $H$ are evaluated at the resident dimorphic point $(\vec{z}_u, \vec{z}_v, p_*)$. Again, the eigenvalues are the diagonal elements, and they determine the definteness of $\mathbf{H}$. The same is not true for $\mathbf{J}$ below.

To define the Jacobian, we refer to equation (10) in [Leimar, 2009], which defines the matrix $\mathbf{Q}$ of cross-derivatives of fitness with respect to all mutant traits and all resident traits. We can denote a block of such cross derivatives where the mutant is from type $k$ and the resident is from type $l$ as follows:

$$(\mathbf{Q}_{kl})_{ij} = \frac{\partial^2 w}{\partial z_{\text{mut},i} \partial z_{l,j}} \Big|_{\vec{z}_{\text{mut}}=\vec{z}_k}$$

The desired Jacobian matrix is evaluated $\mathbf{J} = \mathbf{H}+\mathbf{Q}$. We write this $4 \times 4$ matrix below in two blocks: the left half ($4 \times 2$) and the right half ($4 \times 2$).

$\mathbf{J} = \mathbf{H} + \mathbf{Q} =$

$$\text{Left:} \begin{bmatrix} \left(\frac{\partial^2 w}{\partial z_{\text{mut},1}^2} + \frac{\partial^2 w}{\partial z_{\text{mut},1} \partial z_{u,1}}\right)\Big|_{z_{\text{mut},1}=z_{u,1}} & 0 \\ 0 & \left(\frac{\partial^2 w}{\partial z_{\text{mut},2}^2} + \frac{\partial^2 w}{\partial z_{\text{mut},2} \partial z_{u,2}}\right)\Big|_{z_{\text{mut},2}=z_{u,2}} \\ \frac{\partial^2 w}{\partial z_{\text{mut},1} \partial z_{u,1}}\Big|_{z_{\text{mut},1}=z_{v,1}} & 0 \\ 0 & \frac{\partial^2 w}{\partial z_{\text{mut},2} \partial z_{u,2}}\Big|_{z_{\text{mut},2}=z_{v,2}} \end{bmatrix}$$

$$\text{Right:} \begin{bmatrix} \frac{\partial^2 w}{\partial z_{\text{mut},1} \partial z_{v,1}}\Big|_{z_{\text{mut},1}=z_{u,1}} & 0 \\ 0 & \frac{\partial^2 w}{\partial z_{\text{mut},2} \partial z_{v,2}}\Big|_{z_{\text{mut},2}=z_{u,2}} \\ \left(\frac{\partial^2 w}{\partial z_{\text{mut},1}^2} + \frac{\partial^2 w}{\partial z_{\text{mut},1} \partial z_{v,1}}\right)\Big|_{z_{\text{mut},1}=z_{v,1}} & 0 \\ 0 & \left(\frac{\partial^2 w}{\partial z_{\text{mut},2}^2} + \frac{\partial^2 w}{\partial z_{\text{mut},2} \partial z_{v,2}}\right)\Big|_{z_{\text{mut},2}=z_{v,2}} \end{bmatrix} \tag{51}$$

### D Analysis with payoffs of the form $\pi = B \cdot C$

#### D.1 Fitness

The fitness $w$ of a mutant with trait $z_{\text{mut}}$ in a population with resident trait $z_{\text{res}}$ is

$$w(z_{\text{mut}}, z_{\text{res}}, r) = C(z_{\text{mut}}) \cdot \sum_{k=0}^{n-1} \binom{n-1}{k} r^k (1-r)^{n-1-k} B(\vec{z}_{k+1}) \tag{52}$$

$$\text{where} \qquad \vec{z}_{k+1} = \big(\underbrace{z_{\text{mut}}, \ldots, z_{\text{mut}}}_{k+1}, \underbrace{z_{\text{res}}, \ldots, z_{\text{res}}}_{n-k-1}\big) \tag{53}$$

That is, $\vec{z}_{k+1} := (z_1, z_2, \ldots, z_n)$ st. $z_i = z_{\text{mut}}$ $\forall i \leq k+1$ and $z_i = z_{\text{res}}$ $\forall i > k+1$.

Note that

$$\frac{dz_i}{dz_{\text{mut}}} := \begin{cases} 1 & 1 \leq i \leq k+1 \\ 0 & i > k+1 \end{cases} \tag{54}$$

and $\frac{dz_j}{dz_{\text{mut}}}$ is defined analogously.

We assume that $\frac{dC}{dz_i} < 0$ and $\frac{\partial B}{\partial z_i} > 0$ for any $i \in \{1, ..., n\}$.

#### D.2 Convergence stability

We derive the selection gradient as follows:

$$\frac{\partial w}{\partial z_{\text{mut}}} = \left(C(z_{\text{mut}}) \sum_{k=0}^{n-1} \binom{n-1}{k} r^k (1-r)^{n-1-k} \Big(\sum_{i=1}^{n} \frac{\partial B}{\partial z_i} \frac{dz_i}{dz_{\text{mut}}}\Big)\right) + \left(\frac{dC}{dz_{\text{mut}}} \sum_{k=0}^{n-1} \binom{n-1}{k} r^k (1-r)^{n-1-k} B(\vec{z}_{k+1})\right) \tag{55}$$

$$\frac{\partial w}{\partial z_{\text{mut}}}\Big|_{z_{\text{mut}}=z_{\text{res}}=z} = C(z)\sum_{k=0}^{n-1}\binom{n-1}{k} r^k(1-r)^{n-1-k}(k+1)\frac{\partial B}{\partial z} + \frac{dC}{dz}B(\vec{z})$$
$$= C(z)(1+(n-1)r)B_\bullet + C_\bullet B(\vec{z}) \quad (56)$$

Where $C_\bullet = \frac{dC}{dz_\bullet}$ and $B_\bullet = \frac{\partial B}{\partial z_\bullet}$ are the derivatives of $C$ and $B$ wrt. a focal investment ($\bullet$) among the $n$, when all are evaluated at the resident point $z$. To derive equation (4), we assume exchangeability of investments evaluated at the same point.

The CS condition (inequality 14 in WL (2014)) is

$$\frac{d}{dz}\left[C(z)(1+(n-1)r)B_\bullet + C_\bullet B(\vec{z})\right] < 0$$
$$C(z)(1+(n-1)r)(B_{\bullet,\bullet} + (n-1)B_{\bullet,\bullet'}) + C_\bullet(1+(n-1)r)B_\bullet + C_\bullet(nB_\bullet) + C_{\bullet,\bullet}B(\vec{z}) < 0 \quad (57)$$

Where $C_{\bullet,\bullet} = \frac{d^2C}{dz^2}$ is the second derivative of the cost function, $B_{\bullet,\bullet} = \frac{\partial^2 B}{\partial z_\bullet^2}\big|_{z_\bullet=z\ \forall\bullet}$ is the second derivative of the benefit function wrt. any focal trait, and $B_{\bullet,\bullet'} = \frac{\partial^2 B}{\partial z_\bullet \partial z_{\bullet'}}\big|_{z_\bullet=z_{\bullet'}=z\ \forall\bullet}$ is the cross derivative of the benefit function wrt. focal trait $z_\bullet$ and any nonfocal trait $z_{\bullet'}$, where all are evaluated at resident trait $z$.

#### D.3 Evolutionary instability

Return to equation (3) above and take the second partial derivative of fitness wrt. $z_{\text{mut}}$:

$$\frac{\partial^2 w}{\partial z_{\text{mut}}^2} = C(z)\sum_{k=0}^{n-1}\binom{n-1}{k} r^k(1-r)^{n-1-k}\Big[\sum_{j=1}^{n}\frac{\partial}{\partial z_j}\Big(\sum_{i=1}^{n}\frac{\partial B}{\partial z_i}\frac{dz_i}{dz_{\text{mut}}}\Big)\frac{dz_j}{dz_{\text{mut}}}\Big]$$
$$+ 2\frac{\partial C}{\partial z_{\text{mut}}}\sum_{k=1}^{n-1} r^k(1-r)^{n-1-k}\sum_{i=1}^{n}\frac{\partial B}{\partial z_i}\frac{dz_i}{dz_{\text{mut}}}$$
$$+ \frac{d^2C}{dz_{\text{mut}}^2}\sum_{k=0}^{n-1}\binom{n-1}{k} r^k(1-r)^{n-1-k}B(\vec{z}_{k+1}) \quad (58)$$

Now evaluate at a resident trait, such that $z_{\text{mut}} = z_{\text{res}} = z$:

$$\frac{\partial^2 w}{\partial z_{\text{mut}}^2}\Big|_{z_{\text{mut}}=z_{\text{res}}=z} = C(z)\Big((1+(n-1)r)B_{\bullet,\bullet} + (n-1)r(2+(n-2)r)B_{\bullet,\bullet'}\Big) + 2C_\bullet(1+(n-1)r)B_\bullet + C_{\bullet,\bullet}B(\vec{z}) \quad (59)$$

#### D.4 Branching requires that the cross derivative is negative ($B_{\bullet,\bullet'} < 0$)

From (6), we have

$$C(z)(1+(n-1)r)(n-1)B_{\bullet,\bullet'} + nC_\bullet B_\bullet < -\big(C(z)(1+(n-1)r)B_{\bullet,\bullet} + C_\bullet(1+(n-1)r)B_\bullet + C_{\bullet,\bullet}B(\vec{z})\big)$$

From (8), we have

$$C(z)(n-1)r(2+(n-2)r)B_{\bullet,\bullet'} + C_\bullet(1+(n-1)r)B_\bullet > -\big(C(z)(1+(n-1)r)B_{\bullet,\bullet} + C_\bullet(1+(n-1)r)B_\bullet + C_{\bullet,\bullet}B(\vec{z})\big)$$

Taking the leftmost and rightmost terms, we have

$$C(z)(1+(n-1)r)(n-1)B_{\bullet,\bullet'} + nB_{\bullet}C_{\bullet} \; < \; C(z)(n-1)r(2+(n-2)r)B_{\bullet,\bullet'} + (1+(n-1)r)B_{\bullet}C_{\bullet} \qquad (60)$$

This implies $B_{\bullet,\bullet'} < 0$ for branching, by the same logic as with $\pi = B - C$.

### E Proving properties of interchangeability

For clarity, write $B = B(z_i(x,y), z_j(x,y))$. From definition 1, we have:

$$\begin{aligned}
\frac{\partial}{\partial x} B(\vec{z})\Big|_{z_i=x,\, z_j=y} &= \frac{\partial}{\partial x} B(\vec{z})\Big|_{z_i=y,\, z_j=x} \\
\left(\frac{\partial B}{\partial z_i}\frac{\partial z_i}{\partial x} + \frac{\partial B}{\partial z_j}\frac{\partial z_j}{\partial x}\right)\Big|_{z_i=x,\, z_j=y} &= \left(\frac{\partial B}{\partial z_i}\frac{\partial z_i}{\partial x} + \frac{\partial B}{\partial z_j}\frac{\partial z_j}{\partial x}\right)\Big|_{z_i=y,\, z_j=x} \\
\left(\frac{\partial B}{\partial z_i}(1) + \frac{\partial B}{\partial z_j}(0)\right)\Big|_{z_i=x,\, z_j=y} &= \left(\frac{\partial B}{\partial z_i}(0) + \frac{\partial B}{\partial z_j}(1)\right)\Big|_{z_i=y,\, z_j=x} \\
\frac{\partial B}{\partial z_i}\Big|_{z_i=x,\, z_j=y} &= \frac{\partial B}{\partial z_j}\Big|_{z_i=y,\, z_j=x} \qquad \text{(Property 1)}
\end{aligned}$$

---

$$\begin{aligned}
\frac{\partial}{\partial x}\frac{\partial B}{\partial z_i}\Big|_{z_i=x,\, z_j=y} &= \frac{\partial}{\partial x}\frac{\partial B}{\partial z_j}\Big|_{z_i=y,\, z_j=x} \\
\left(\frac{\partial^2 B}{\partial z_i^2}\frac{\partial z_i}{\partial x} + \frac{\partial^2 B}{\partial z_i \partial z_j}\frac{\partial z_j}{\partial x}\right)\Big|_{z_i=x,\, z_j=y} &= \left(\frac{\partial^2 B}{\partial z_j \partial z_i}\frac{\partial z_i}{\partial x} + \frac{\partial^2 B}{\partial z_j^2}\frac{\partial z_j}{\partial x}\right)\Big|_{z_i=y,\, z_j=x} \\
\left(\frac{\partial^2 B}{\partial z_i^2}(1) + \frac{\partial^2 B}{\partial z_i \partial z_j}(0)\right)\Big|_{z_i=x,\, z_j=y} &= \left(\frac{\partial^2 B}{\partial z_j \partial z_i}(0) + \frac{\partial^2 B}{\partial z_j^2}(1)\right)\Big|_{z_i=y,\, z_j=x} \\
\frac{\partial^2 B}{\partial z_i^2}\Big|_{z_i=x,\, z_j=y} &= \frac{\partial^2 B}{\partial z_j^2}\Big|_{z_i=y,\, z_j=x} \qquad \text{(Property 2)}
\end{aligned}$$

---

$$\begin{aligned}
\frac{\partial}{\partial y}\frac{\partial B}{\partial z_i}\Big|_{z_i=x, z_j=y} &= \frac{\partial}{\partial y}\frac{\partial B}{\partial z_j}\Big|_{z_i=y, z_j=x} \\
\left(\frac{\partial^2 B}{\partial z_i^2}\frac{\partial z_i}{\partial y} + \frac{\partial^2 B}{\partial z_i \partial z_j}\frac{\partial z_j}{\partial y}\right)\Big|_{z_i=x,\, z_j=y} &= \left(\frac{\partial^2 B}{\partial z_j \partial z_i}\frac{\partial z_i}{\partial y} + \frac{\partial^2 B}{\partial z_j^2}\frac{\partial z_j}{\partial y}\right)\Big|_{z_i=y,\, z_j=x} \\
\left(\frac{\partial^2 B}{\partial z_i^2}(0) + \frac{\partial^2 B}{\partial z_i \partial z_j}(1)\right)\Big|_{z_i=x,\, z_j=y} &= \left(\frac{\partial^2 B}{\partial z_j \partial z_i}(1) + \frac{\partial^2 B}{\partial z_j^2}(0)\right)\Big|_{z_i=y,\, z_j=x} \\
\frac{\partial^2 B}{\partial z_i \partial z_j}\Big|_{z_i=x, z_j=y} &= \frac{\partial^2 B}{\partial z_i \partial z_j}\Big|_{z_i=y, z_j=x} \qquad \text{(Property 3)}
\end{aligned}$$

Properties 1-2 hold if the indices $i$ and $j$ are switched. This can be shown by applying the same derivation as above, taking $\frac{\partial}{\partial y}$ for property 1, then $\frac{\partial^2}{\partial y^2}$ for property 2. Property 3 does not change if the indices are switched.

### F Proving uniqueness and dynamical stability of $p_*$

**Result** *Let the payoff function for one public good for player $i \in (1, ..., n)$ be $\pi(z_i, \cdot) = B(g(z_i, \cdot)) - C(z_i)$ where the function $g(z_i, \cdot) = \sum_{j=1}^{n} z_j$ (with other assumptions as defined in Methods), and let the investments of dimorphic* *types be (wlog)* $z_u > z_v \geq 0$.

*If* $B'' := \frac{d^2 B}{dg^2} < 0$ *on* $\mathbb{R}^+$ *and interior solution* $p_* \in (0,1)$ *satisfying* $\dot{p} = 0$ *exists, then* $p_*$ *is unique and stable under replicator dynamics.*

**Proof:** We apply Result 3 of Peña et al. [2014a], which the authors extended in Peña et al. [2014b] to incorporate relatedness among individuals.

The focal individual plays with $n-1$ others. Let $a_q$ be the payoff to an individual investing $z_u$ when $q \leq n-1$ others invest $z_u$ (and $n-1-q$ players invest $z_v$), and let $b_q$ be the payoff to an individual investing $z_v$ when $q$ others invest $z_u$. Define $d_q := a_q - b_q$ and denote $\triangle d_q = d_{q+1} - d_q$. Define $e_q := q\triangle a_{q-1} + (n-1-q)\triangle b_q$, where $\triangle a_{q-1} = a_q - a_{q-1}$ and $\triangle b_q = b_{q+1} - b_q$. Finally, define $f_q := d_q + re_q$. Denote $\triangle f_q = f_{q+1} - f_q$ and denote the gain sequence $\mathbf{f} := (f_0, f_1, ..., f_{n-1})$.

These definitions follow Peña et al. [2014b] section 2.4, where we have used the subscript $q$ instead of the authors' $k$ to distinguish from the subscript $k$ (number of relatives) used in our definition of fitness. Under our simple relatedness scheme, the scaled relatedness coefficient $\kappa$ from Peña et al. [2014b] is equal to the parameter $r$.

First, we expand $d_q$ and $e_q$ and derive $f_q$ using the definition of payoff above:

$$\begin{aligned}
a_q &= B\big(z_u + qz_u + (n-1-q)z_v\big) - C(z_u) \\
a_{q-1} &= B\big(z_u + (q-1)z_u + (n-q)z_v\big) - C(z_u) \\
b_q &= B\big(z_v + qz_u + (n-1-q)z_v\big) - C(z_v) \\
b_{q+1} &= B\big(z_v + (q+1)z_u + (n-2-q)z_v\big) - C(z_v) \\
\triangle a_{q-1} = a_q - a_{q-1} &= B\big((q+1)z_u + (n-1-q)z_v\big) - B\big(qz_u + (n-q)z_v\big) \\
\triangle b_q = b_{q+1} - b_q &= B\big((q+1)z_u + (n-1-q)z_v\big) - B\big(qz_u + (n-q)z_v\big)
\end{aligned}$$

$$\begin{aligned}
d_q &= a_q - b_q = B\big(z_u + qz_u + (n-1-q)z_v\big) - C(z_u) - \Big(B\big(z_v + qz_u + (n-1-q)z_v\big) - C(z_v)\Big) \\
&= \Big(B\big((q+1)z_u + (n-1-q)z_v\big) - B\big(qz_u + (n-q)z_v\big)\Big) - \Big(C(z_u) - C(z_v)\Big) \\
e_q &= q\triangle a_{q-1} + (n-1-q)\triangle b_q \\
&= (n-1)\Big(B\big((q+1)z_u + (n-1-q)z_v\big) - B\big(qz_u + (n-q)z_v\big)\Big)
\end{aligned}$$

$$f_q = d_q + re_q = (1+(n-1)r)\Big(B\big((q+1)z_u + (n-1-q)z_v\big) - B\big(qz_u + (n-q)z_v\big)\Big) - \Big(C(z_u) - C(z_v)\Big) \quad (61)$$

Following Peña et al. [2014b] equation (10), we can now write the gain function $G$:

$$G(p) = \sum_{q=0}^{n-1} \binom{n-1}{q} p^q (1-p)^{n-1-q} f_q \quad (62)$$

By Result 3 of Peña et al. [2014a], if the gain sequence $\mathbf{f}$ has a single sign change and $f_0 > 0$, then any interior solution $p_* \in (0,1)$ satisfying $\dot{p} = 0$ is unique and stable under replicator dynamics.

Note that we have assumed that public benefit $B$ is a function of the sum of investments, denoting $g(z_i, \cdot) = \sum_{j=1}^{n} z_j$. We now show that if $\frac{d^2 B}{dg^2} < 0$ on $\mathbb{R}^+$, then $\triangle f_q < 0$.

$$\begin{aligned}
\triangle f_q &= f_{q+1} - f_q \\
&= (1+(n-1)r)B\big((q+2)z_u + (n-2-q)z_v\big) - C(z_u) - \Big((1+(n-1)r)B\big((q+1)z_u + (n-1-q)z_v\big) - C(z_v)\Big) \\
&\quad - \Bigg((1+(n-1)r)B\big((q+1)z_u + (n-1-q)z_v\big) - C(z_u) - \Big((1+(n-1)r)B\big(qz_u + (n-q)z_v\big) - C(z_v)\Big)\Bigg) \\
&= (1+(n-1)r)\Big[B\big((q+2)z_u + (n-2-q)z_v\big) - B\big((q+1)z_u + (n-1-q)z_v\big) \\
&\quad - \Big(B\big((q+1)z_u + (n-1-q)z_v\big) - B\big(qz_u + (n-q)z_v\big)\Big)\Big] \\
&= (1+(n-1)r)\Big[B\big(2(z_u - z_v) + qz_u + (n-q)z_v\big) - B\big((z_u - z_v) + qz_u + (n-q)z_v\big) \\
&\quad - \Big(B\big((z_u - z_v) + qz_u + (n-q)z_v\big) - B\big(qz_u + (n-q)z_v\big)\Big)\Big]
\end{aligned}$$

If $B'' < 0$, the term in square brackets in the last line above is negative. This is because for any continuous, real-valued function $h(x)$, for $x_0 < x_1 < x_2$ with $x_1 - x_0 = x_2 - x_1$, $\frac{\partial^2 h}{\partial x^2} < 0$ implies that $h(x_2) - h(x_1) < h(x_1) - h(x_0)$. Also, the term $(1+(n-1)r)$ is strictly positive. We conclude that $\triangle f_q < 0$.

Lastly, to show that the gain sequence $\mathbf{f}$ has a single sign change given $\triangle f_q < 0$, we must show that $f_0 > 0$ and $f_{n-1} < 0$. Rewrite both of these expressions as follows:

$$\begin{aligned}
f_0 > 0 &\Rightarrow (1+(n-1)r)\Big(B\big(z_u + (n-1)z_v\big) - B\big(nz_v\big)\Big) - \Big(C(z_u) - C(z_v)\Big) > 0 \\
&\Rightarrow (1+(n-1)r)\Big(B\big(z_u + (n-1)z_v\big) - B\big(nz_v\big)\Big) > C(z_u) - C(z_v)
\end{aligned}$$

$$\begin{aligned}
f_{n-1} < 0 &\Rightarrow (1+(n-1)r)\Big(B\big(nz_u\big) - B\big((n-1)z_u + z_v\big)\Big) - \Big(C(z_u) - C(z_v)\Big) > 0 \\
&\Rightarrow (1+(n-1)r)\Big(B\big(nz_u\big) - B\big((n-1)z_u + z_v\big)\Big) > C(z_u) - C(z_v)
\end{aligned}$$

At a candidate solution $p_* \in (0,1)$ for which $\dot{p} = 0$, the following equality is true (see equations (40) and (41)):

$$w(z_u, z_u, z_v, p, r) = w(z_v, z_u, z_v, p, r)$$

$$-C(z_u) + \sum_{k=0}^{n-1} \sum_{l=0}^{n-1-k-l} \frac{(n-1)!}{k!\, l!(n-1-k-l)!} r^k \big((1-r)p\big)^l \big((1-r)(1-p)\big)^{n-1-k-l} B(z_u + (k+l)z_u + (n-1-k-l)z_v)$$

$$= -C(z_v) + \sum_{k=0}^{n-1} \sum_{l=0}^{n-1-k-l} \frac{(n-1)!}{k!\, l!(n-1-k-l)!} r^k \big((1-r)p\big)^l \big((1-r)(1-p)\big)^{n-1-k-l} B(z_v + (k+l)z_u + (n-1-k-l)z_v)$$

$$\sum_{k=0}^{n-1} \sum_{l=0}^{n-1-k-l} \frac{(n-1)!}{k!\, l!(n-1-k-l)!} r^k \big((1-r)p\big)^l \big((1-r)(1-p)\big)^{n-1-k-l} \Big(B(z_u, \cdot) - B(z_v, \cdot)\Big) = C(z_u) - C(z_v)$$

$$\mathbb{E}\Big[B(z_u + (k+l)z_u + (n-1-k-l)z_v) - B(z_v + (k+l)z_u + (n-1-k-l)z_v)\Big] = C(z_u) - C(z_v) \tag{63}$$

Show by contradiction that $B'' < 0$ implies $f_0 > 0$. Assume that $f_0 < 0$. We have

$$(1 + (n-1)r)\Big(B\big(z_u + (n-1)z_v\big) - B\big(nz_v\big)\Big) - \Big(C(z_u) - C(z_v)\Big) < 0$$

$$(1 + (n-1)r)\Big(B\big(z_u + (n-1)z_v\big) - B\big(nz_v\big)\Big) < \Big(C(z_u) - C(z_v)\Big)$$

Because $B'' < 0$, every value of $B$ in the LHS expectation in equation (63) (i.e. with $q > 0$) is smaller than the interval $B\big(z_u + (n-1)z_v\big) - B(nz_v)$. This implies

$$\mathbb{E}\Big[B(z_u{+}qz_u{+}(n{-}1{-}q)z_v){-}B(z_v{+}qz_u{+}(n{-}1{-}q)z_v)\Big] \leq (1{+}(n{-}1)r)\Big(B\big(z_u{+}(n{-}1)z_v\big){-}B\big(nz_v\big)\Big) < \Big(C(z_u){-}C(z_v)\Big),$$

where $q = k + l$. This contradicts equation (63), so we conclude $f_0 > 0$.

Similarly, assume that $f_{n-1} > 0$. We have

$$(1 + (n-1)r)\Big(B\big(nz_u\big) - B\big((n-1)z_u + z_v\big)\Big) > \Big(C(z_u) - C(z_v)\Big)$$

Because $B'' < 0$, every value of $B$ in the LHS expectation in equation (63) is larger than the interval $B\big(nz_u\big) - B\big((n-1)z_u + z_v\big)$. This implies

$$\mathbb{E}\Big[B(z_u{+}qz_u{+}(n{-}1{-}q)z_v){-}B(z_v{+}qz_u{+}(n{-}1{-}q)z_v)\Big] \geq (1{+}(n{-}1)r)\Big(B\big(nz_u\big){-}B\big((n{-}1)z_u{+}z_v\big)\Big) > \Big(C(z_u){-}C(z_v)\Big),$$

This again contradicts equation (63), so we conclude $f_{n-1} < 0$. ■
