## Supplementary code for "The effects of group size and assortment on the evolution of division of labor": AD-withr-BDprocess-code.py

```
import pandas as pd 
import numpy as np
import math
import matplotlib.pyplot as plt
import numpy.random as rnd
import pickle
import time
import json
from copy import deepcopy as dp
from math import comb, exp
from scipy.stats import multinomial, binom
from itertools import chain, repeat
import itertools

# DEFINE KEY FUNCTIONS

# this function returns a list of all possible phenotypic configurations of the nonfocal, nonrelated individuals in a focal’s PG group, given the number of phenotypic types in the population. Note that gsize <= n-1 here.
def get_multinomial_support(gsize,ntypes):
    if ntypes==1:
        return np.array([[gsize]])
    arr = np.arange(0,gsize)
    st = ('arr,')*(ntypes-2) + 'arr'
    bigmat = eval('np.meshgrid('+st+')')
    xkarr = gsize - bigmat[0].flatten()
    for k in range(1,ntypes-1):
        xkarr = xkarr - bigmat[k].flatten()
    mat = np.array([bigmat[k].flatten() for k in range(ntypes-1)] + [xkarr]).T
    mat = np.array([g for g in mat if g[ntypes-1] >= 0])
    for k in range(ntypes-1):
        mat=np.concatenate((mat,np.array([[gsize if j==k else 0 for j in range(ntypes-1)]+[0]])),axis=0)
    return mat 


# function to calculate the (multinomial) probability of a given phenotypic configuration of the nonfocal nonrelated individuals in a focal’s PG group
def get_multinomial_prob(gsize,config,frqs): 
    prod = 1
    for ff in config:
        prod *= math.factorial(ff)
    term1 = math.factorial(gsize)/prod
    for i in range(numtypes):
        term1 *= frqs[i]**(config[i])
    return term1


# binary search function for use use when choosing individuals to give birth proportional to their fitness
def binary_search(cutoffs,s,low,high):
    mid=int(np.floor((low+high)/2))
    ntypes = len(cutoffs)
    if s<=cutoffs[0]:
        return 0
    elif s>cutoffs[ntypes-2]:
        return ntypes-1
    elif (s>cutoffs[mid-1] and s<=cutoffs[mid]):
        return mid
    elif s>cutoffs[mid]:
        return binary_search(cutoffs,s,mid+1,high)
    elif s<cutoffs[mid]:
        return binary_search(cutoffs,s,low,mid-1)

# function to calc model 1 payoff for focal trait z and list of n traits "grp”. The list of length n includes focal, related, and nonrelated nonfocal group members
def payoff(z,grp):
    return (sum(grp))**a1 - z**a2


# function to calc expected payoffs for all types in current pop state, given by traits and frequencies. Refer to equation 7(?) in manuscript.
def calcfitnesses(trts,frqs):  
    return [exp(s * (r**(n-1) * payoff(typ, [typ for i in range(n)]) + sum([binom.pmf(numrels,n-1,r) * 
        sum([get_multinomial_prob(n - 1 - numrels, g, freqs) * payoff(typ, [typ for i in range(numrels+1)] + list(chain.from_iterable(repeat(trait, freq) for trait, freq in zip(trts, g)))) 
            for g in configs_list[(n - 1 - numrels,numtypes)]]) for numrels in range(n-1)]) ) )  
            for typ in trts]

# this function returns a “cutoff” list of which each element i<=numtypes is the cumulative sum of the fitness (weighted by frequencies and normalized ) of all types <= i.  For use use when choosing individuals to give birth proportional to their fitness.
def get_birth_cutoffs(fits,frqs):  #fitnesses,numtypes,numpertype
    weighted = [frqs[i]*fits[i] for i in range(numtypes)]
    normd = [i/sum(weighted) for i in weighted]
    return np.cumsum(normd)   

# this function returns a cutoff list where cutoff size is only proportional to frequency.  For use use when choosing individuals to die
def get_death_cutoffs(frqs):  #numtypes,numpertype,N
    ct = 0
    cutoffs = [0 for i in range(numtypes)]
    for i in range(numtypes):
        ct += frqs[i]
        cutoffs[i] = ct
    return cutoffs


# INITIALIZE

# params
N= 1000   # finite population size
step = 1/N  
n = 4 #size of each PG game group in a round
r = 0   # relatedness
s = 100   # selection strength (affecting wall time of sim)

a1 = .7    # governs concavity of public benefit function 
a2 = .8   # governs concavity of private cost function

mutprob = .0001     # probability of mutation per birth
mutvar = .01     # standard deviation of normally distributed mutation

zcrit = ( (1 + (n - 1)*r) * n**(a1 - 1) * a1/a2 )**(1/(a2 - a1))    # compute critical point, for reference in deciding initial pop state
print('branching conditions ',(1/n) + ((n - 1)/n) * (r * (2 + (n - 2)* r)/(1 + (n - 1)* r)) < (a2 - 1)/(a1 - 1) < 1 )   # if true, branching conditions are met

#simulation parameters 
runs = 1    # number of runs of the simulation with the params above 
tsteps = int(.1*10**7)+1    # number of timesteps in one run of sim
trecord = int(10**4)     # tstep interval at which population state is recorded in ‘poprec’
tsave = int(10**6)     # tstep interval at which poprec is written to disk as a json
tprint = int(.1*10**6)     # tstep interval at which current pop info prints 

poprec = {'params':{'N':N,'n':n,'r':r,'mutprob':mutprob,'mutsize':mutvar,'s':s,'tinterval':trecord,'numsteps':tsteps,'model':
                    {'a1':a1,'a2':a2}}}

skip = False   # when the type that gives birth is the same as the type that dies and there is no mutation, the remainder of that timestep can be skipped. 

# Pre-populate dict of lists of phenotypic configurations of nonfocal, nonrelated group members, for subgroup sizes ranging from 1 to n-1, and number of types up to maxnumtypes.
configs_list={}
maxnumtypes = 9  # increment up as needed during sim
for gsize in range(1,n):
    for ntypes in range(1,maxnumtypes+1):
        configs_list[(gsize,ntypes)] = get_multinomial_support(gsize,ntypes)

# RUN SIM:

if __name__ == '__main__': # to run in terminal, ensure everything below is indented

    for run in range(runs):
        # initialize population :
        numtypes=1   # number of existing types at t=0 (updated over time)
        traits = [.1]     # list of existing traits at t=0    (updated over time)
        freqs = [1]     # list of frequencies of existing types, with index corresponding to ‘traits’  (updated over time)
        fitnesses = calcfitnesses(traits,freqs)      # list of fitnesses of existing types, with index corresponding to ‘traits’ and ‘freqs’ (updated over time)
        poprec[run]={}     # initialize subdictionary corresponding to this run in the global poprec dictionary
        print('run: ',run)
        t0=time.time()
        skip=False

        for t in range(tsteps):
            if t%trecord == 0:   # at interval trecord, record current traits, freqs, and fitnesses of existing types
                poprec[run][t] = dp({traits[i]:freqs[i] for i in range(numtypes)})
                poprec[run][t]['fitnesses']=fitnesses
        
            if t%tsave==0:    # save poprec to disk as json
                json_obj=json.dumps(poprec)
                with open('filename-branchingsim','w') as outfile:   
                    outfile.write(json_obj)
                
            if t%tprint==0:   # printing valuable info at interval tprint
                print('t: ',t)
                print(traits,freqs)
                print(fitnesses)
                if t>0: print(b_cutoffs)
                #print('time rate/5M: ', times[len(times)-1])
                tf=time.time()   # wall time rate of sim
                print('wall time per ',tprint,' timesteps: ',tf-t0)
                t0=time.time()

            if skip is False:    # if NOT same type both gave birth and died with no mutations:
                b_cutoffs = get_birth_cutoffs(fitnesses,freqs)     # compute cutoff list for births (defined in functions) from current fitness and frequencies in pop
                d_cutoffs = np.cumsum(freqs) #get_death_cutoffs(freqs)  # compute cutoff list for deaths (defined in functions) from current frequencies in pop

            # get type that will reproduce (proportional to fitness):
            seln_B=rnd.uniform()
            k = binary_search(b_cutoffs, seln_B, 0, numtypes)

            # get type that will die (uniform prob):
            seln_D = rnd.uniform()
            d = binary_search(d_cutoffs, seln_D, 0, numtypes)

            if rnd.uniform() < mutprob: # with low prob, offspring mutates:
                parz = traits[k]
                zmut = parz + rnd.normal(0,mutvar)   # + rnd.uniform(-mutvar,mutvar)   # the commented out option draws mutants from a uniform distribution centered at parent trait
                while zmut <= 0:    # if a drawn mutant trait drops below 0 (not defined), draw again (leading to truncated (at 0) normal distribution)
                    zmut = parz + rnd.normal(0,mutvar)
            
                # Update tickers and records given mutation
                numtypes+=1        # one new type
                traits.append(zmut)  # add mutant trait to traits list
                freqs[d] -= step    # increment down the frequency of the type of the individual selected to die
                freqs.append(step)   # add the frequency of the mutant, equal to step = 1/N
            
                if numtypes>maxnumtypes:      #  if the number of existing types is greater than the max number for which the multinomial support has been precomputed:
	        # compute the new multinomial support with the current number of types
                    for gsize in range(1,n):
                        configs_list[(gsize,numtypes)] = get_multinomial_support(gsize,numtypes)
                    maxnumtypes+=1    # increment maxnumtypes
                
                fitnesses = calcfitnesses(traits,freqs)      # recompute all fitnesses including the mutant’s
                skip = False      # we will recompute birth and death cutoff lists in next timestep
            else:
                if k != d:     # if the type selected for a birth (k) is different than the type selected to a death (d):
                    freqs[k] += step   # increment up freq of birth type by 1/N
                    freqs[d] -= step    # increment down freq of death type by 1/N
                    fitnesses = calcfitnesses(traits,freqs)      # recompute fitness of all types with new frequencies
                    skip=False    # we will recompute birth and death cutoff lists in next timestep
                else:
                    skip = True     # we will not recompute birth and death cutoff lists in next timestep
                    continue      # skip extinction step

            if freqs[d] <= 0:   # if the frequency of a type is equal to 0:
	    #delete the type from state variables
                del freqs[d]
                del traits[d]
                numtypes -= 1      # subtract 1 from the number of existing types
                fitnesses = calcfitnesses(traits,freqs)     # recompute fitness of all types   
                skip=False     # we will recompute birth and death cutoff lists in next timestep


    # PLOTTING:
    # Preprocess the record dictionary, which includes phenotypes, frequencies, and their fitnesses for each timestep
    popdict = {}
    pop=poprec[0]
    fitlist=[]

    for t, subdict in poprec[0].items():
        # Initialize empty subdicts
        popdict[t] = {}
    
        for key, value in subdict.items():
            if key == 'fitnesses':
                # Put fitness list into the fitnesses dict
                fitlist=fitlist+value
            else:
                # Trait keys (float) go into the traits dict
                popdict[t][key] = value

    # the following code plots the evolutionary trajectory for a given run:
    # at a timepoint (given in generation-time) on the horizontal axis, the traits of existing types are plotted vertically as dots, with their opacity weighted by the frequency of the existing types at that timepoint
    # The monomorphic critical point zcrit is plotted as a horizontal dotted line.
    # note this code uses popdict from working memory (ie not converting strings).

    x=list(itertools.chain.from_iterable([[t/N for k in range(len(popdict[t]))] for t in popdict]))      # list of timepoints (scaled by generation time = N timesteps), each with multiplicity equal to the number of existing types at that timepoint
    y=list(itertools.chain.from_iterable([k.keys() for k in popdict.values()]))         # flattened list of all traits existing at every timepoint (with corresponding time markers given by x)

    fittest=max(fitlist)      # maximum fitness of all types throughout run
    relfits = [i/fittest for i in fitlist]       # all fitnesses normalized to maximum
    colors = [(0, 0, 0, alpha**.11) for alpha in relfits]      # list of colors, with opacity weighted by fitness of type at given timepoint. The lower the exponent of alpha, the faster opacity increases with fitness

    #plot data using matplotlib:
    tmin=0
    tmax=int(len(x))
    cargs={'fontname':'helvetica'}
    plt.scatter(x[tmin:tmax],y[tmin:tmax],c=colors[tmin:tmax])
    plt.xlabel('Generation',{'fontname':'helvetica'})
    plt.ylabel('Investments',{'fontname':'helvetica'})
    plt.xticks(fontsize=16, fontfamily='helvetica')
    plt.yticks(fontsize=16, fontfamily='helvetica')
    plt.axhline(y=zcrit, linestyle='--', color='gray')
    plt.show(block=True)
```
