## Supplementary code for "The effects of group size and assortment on the evolution of division of labor": README-mathematica.txt

This mathematica script is organized into labeled sections. All sections use payoff model 1 for analysis, but can be converted to use payoff model 2.

The first section generates figure 3, which illustrates the fitness of dimorphic mutants (labeled zu and zv) in a monomorphic resident population with critical investment level z*.

The second section generates figure 2. First it imports plots of simulation data generated by the corresponding python script AD-withr-BDprocess-code.py. It plots these next to the branching parameter regions (named "V" in main text).

The third section analyzes the adaptive dynamics of dimorphic types with investments zu and zv after branching, with one public good. For a given parametrization of payoff model 1, it computes the monomorphic critical point z*, initializes zu and zv close to z*, and integrates their two-type adaptive dynamical trajectories. If those trajectories reach a dimorphic critical point, functions Hmat and Jmat can be used to check the convergence and evolutionary (in)stability of said point. 

The fourth section carries out the above procedure for a mesh of parametrizations that allow branching.

The final section analyzes the adaptive dynamics of dimorphic types with investments zu and zv after branching, with two public goods. It generates initial dimorphisms close to a two-dimensional branching point and integrates their adaptive dynamical trajectories. If the latter reach a dimorphic critical point, it checks the stability properties. It also checks the final DOL state of the population (exploitation or specialization, main text), as establishes that it is always equivalent to the initial DOL state.
