## Supplementary code for "The effects of group size and assortment on the evolution of division of labor": README-python.txt

The included python script simulates a standard Moran process in a finite population of size N. Then, for the first run of the simulation, the program plots the traits and relative fitnesses of existing types over generational time.

Fitness of a type is equal to its expected payoff in a Public Goods (PG) game with group size little n, using payoff model 1, with the assortment mechanism described in Methods (main text). 

The program generates a number (equal to parameter 'runs') of instances of the moran process. The parameters for all runs in the program are given in the subsection with comment label #params. The number of timesteps per run can be adjusted in this section. The code is organized into four sections: (1) define functions; (2) initialize (including params); (3) run simulation program (in terminal, this is below the condition "if __name__ == '__main__': ");  (4) plot the phenotypic timeseries for the first run.

The initial state of the population is defined in the timestep 0 (i.e. below "if __name__ == '__main__': "), as the global variables 'traits' and 'freqs', which are lists.

To run the program on terminal: ensure that python3 is installed on your computer; change working directory to wherever this .py script is saved on your computer; and run the command  "python 3 ./AD-withr-BDprocess-code.py" .
